# AIM2 Inflammasome Mediates Hallmark Neuropathological Alterations and Cognitive Impairment in a Mouse Model of Vascular Dementia

**DOI:** 10.1101/2020.06.05.135228

**Authors:** Luting Poh, David Y. Fann, Peiyan Wong, Hong Meng Lim, Sok Lin Foo, Sung-Wook Kang, Vismitha Rajeev, Sharmelee Selvaraji, Vinaya Rajagopal Iyer, Nageiswari Parathy, Mohammad Badruzzaman Khan, David C. Hess, Dong-Gyu Jo, Grant R. Drummond, Christopher G. Sobey, Mitchell K.P. Lai, Christopher Li-Hsian Chen, Lina H. K. Lim, Thiruma V. Arumugam

## Abstract

Chronic cerebral hypoperfusion is associated with vascular dementia (VaD). Cerebral hypoperfusion may initiate complex molecular and cellular inflammatory pathways that contribute to long-term cognitive impairment and memory loss. Here we used a bilateral common carotid artery stenosis (BCAS) mouse model of VaD to investigate its effect on the innate immune response – particularly the inflammasome signaling pathway. Comprehensive analyses revealed that chronic cerebral hypoperfusion induces a complex temporal expression and activation of inflammasome components and their downstream products (IL-1β and IL-18) in different brain regions, and promotes activation of apoptotic and pyroptotic cell death pathways. Polarized glial cell activation, white matter lesion formation and hippocampal neuronal loss also occurred in a spatiotemporal manner. Moreover, in AIM2 knockout mice we observed attenuated inflammasome-mediated production of proinflammatory cytokines, apoptosis and pyroptosis, as well as resistance to chronic microglial activation, myelin breakdown, hippocampal neuronal loss, and behavioural and cognitive deficits following BCAS. Hence, we have demonstrated that activation of the AIM2 inflammasome substantially contributes to the pathophysiology of chronic cerebral hypoperfusion-induced brain injury and may therefore represent a promising therapeutic target for attenuating cognitive impairment in VaD.

## Introduction

Vascular dementia (VaD) is the second most common form of dementia that is associated with vascular diseases resulting in dysfunction and damage to the cerebral vasculature, which causes disruption of cerebral blood flow, brain injury, and ultimately cognitive impairment and memory loss^1^. Clinical findings in VaD include chronic cerebral hypoperfusion, white matter lesions (WMLs), strokes, hippocampal atrophy, decline in executive functioning and memory impairment^2^. Recent evidence indicates that chronic cerebral hypoperfusion is associated with systemic inflammation before WMLs and neurological symptoms develop in VaD^3–5^. Interleukin 1-beta (IL-1β), a potent cytokine that orchestrates inflammatory pathways, is likely to be a key contributor to VaD^5^. The primary molecular machinery for IL-1β production in response involves assembly and activation of the inflammasome.

Inflammasome is an intracellular multi-protein complex that initiates an innate immune response, and is involved in multiple acute and chronic neurological diseases such as ischemic stroke, Alzheimer’s disease (AD), Parkinson’s Disease (PD), and amyotrophic lateral sclerosis (ALS)^6–11^. In the canonical inflammasome pathway, stimulation and homo-oligomerization of inflammasome receptors such as nucleotide-binding domain and leucine-rich repeat containing (NLR) family pyrin domain containing (NLRP) 1 (NLRP1), NLRP3, NLR family CARD domain containing 4 (NLRC4) and absent in melanoma 2 (AIM2) initiates formation of the inflammasome complex, recruitment and activation of adaptor (ASC) and effector (total caspase-1 and -8) inflammasome components into biologically active cleaved caspases-1 and -8^6,12^. In the noncanonical inflammasome pathway, caspase-11 is activated via auto-proteolysis upon dimerization to produce cleaved caspase-11. Upon inflammasome assembly and activation, activity of cleaved caspase-1 and -8 are increased and downstream proinflammatory cytokines such as mature IL-1β and IL-18 are generated^6^. This leads to activation of glial cells, endothelial dysfunction, and oligodendrocyte injury and demyelination, eventually resulting in neurovascular unit disintegration, neuronal loss and brain circuit dysfunction. Cleaved caspase-1 and -8 may also promote cell death by activating apoptosis and pyroptosis pathways^13–14^. Cleaved caspase-1 and -8 induces apoptosis and pyroptosis by catalyzing the cleavage of total caspase-3 and full-length Gasdermin D (GSDMD-FL) into cleaved caspase-3 and N-terminal Gasdermin D (GSDMD-NT), respectively. GSDMD-NT can oligomerize with other N-terminal units and cause pore formation in the nuclear and plasma membranes, leading to leakage of cellular content and proinflammatory cytokines^15,16^. GSDMD activation can also be facilitated directly by non-canonical cleaved caspase-11^17^. The inhibition of caspase-1 was demonstrated to prevent pyroptosis in experimental models of multiple sclerosis by reducing demyelination and neurodegeneration^18^. It was also established that the inflammasome complex also contributed to cognitive decline in a APP/PS1 mouse model of Alzheimer’s disease^9,19^. Specifically, it was shown that deletion of the AIM2 inflammasome in that AD model promoted dendrite branching and synaptic plasticity, and improvement in spatial memory^20^. AIM2 is known to play a role in the host immune response to microbial DNA^21^; however, several reports have established that endogenous double-stranded DNA (dsDNA) can also activate AIM2 to form the AIM2 inflammasome^22,23^. Moreover, AIM2 also responds to dsDNA released from damaged host cells and is a potential ligand that can activate AIM2 in neurons and glial cells following chronic hypoperfusion^22, 24^.

While evidence for direct involvement of the inflammasome complex in VaD is lacking, a cytokine profile of plasma from VaD patients found IL-1β to be the most abundant^25–28^. It was also reported that IL-1β impeded oligodendrocyte recruitment and inhibited white matter repair and functional recovery during the early stages of chronic cerebral hypoperfusion^5^. However, any potential involvement of inflammasome activation in chronic cerebral hypoperfusion-induced glial activation, white matter damage, neuronal cell death and cognitive dysfunction has not been studied.

In the present study, we have comprehensively investigated the role of the inflammasome signaling pathway in the bilateral common carotid artery stenosis (BCAS) mouse model of VaD. The BCAS mouse model has been well established in imitating a wide range of neurological clinical conditions of white matter rarefaction, hippocampal atrophy and cognitive decline induced by chronic cerebral hypoperfusion in humans^29,30^. Moreover, BCAS develops white matter lesions without cerebral infarctions and optic nerve damage, while hippocampal neuronal death and atrophy is induced making it the most appropriate animal model in studying VaD^31^. Using the BCAS mouse model, we report an increased expression and activation of inflammasome complexes, production of mature proinflammatory cytokines, polarized glial activation, WMLs, and neuronal cell death occurring in a time-dependent manner in the cerebral cortex, hippocampus and striatum following BCAS. Notably, these brain regions exhibited distinct profiles of inflammasome expression and activation, and a specific prominent increase of AIM2 in the cerebral cortex and hippocampus. Furthermore, we demonstrated that genetic deletion of AIM2 protects against BCAS-induced inflammasome activation, proinflammatory cytokine production, polarized glial activation, WML formation and neuronal cell death, ultimately leading to an improvement in cognitive outcome. Overall, our findings reveal that the AIM2 inflammasome may be a potential therapeutic target in VaD.

## Materials and Methods

### Experimental Animals and Genotyping

All *in vivo* experimental procedures were approved by the National University of Singapore, Singapore Animal Care and Use Committee and performed according to the guidelines set forth by the National Advisory Committee for Laboratory Animal Research (NACLAR), Singapore. Mice were housed in individual cages under standard laboratory conditions. All efforts were made to minimize suffering and numbers of animals used. All sections of the manuscript were performed and reported in accordance with ARRIVE (Animal Research: Reporting In Vivo Experiments) guidelines. Fourteen to sixteen weeks old wild type (WT) male C57BL/6 mice (weighing 24 to 30 grams) were obtained from In Vivos, Singapore. The AIM2 knockout (B6.129P2-Aim2<Gt(CSG445)Byg>/J; AIM2 KO) mice were generated by Ingenious Targeting Laboratory (Ronkonkoma, NY) on the C57BL/6 background mice via targeted replacement of the AIM2 coding region with a neomycin resistance gene through homologous recombination that was kindly obtained from our collaborator Professor Jenny P. Y. Ting^32^. Primer sequences used for PCR-based AIM2 deficient mouse genotyping are as follows: neomycin (KO) forward, GGAACTTCGCTAGACTAGTACGCGTG; neomycin (KO) reverse, CAACATTGTACAGATTGAGCAGG; AIM2 (WT) forward, GATGGAGAGTGAGTACCGGGAAATGCTGTT; and AIM2 (WT) reverse, TCTGCAAGTAGATTGGAGACAGACTCTGGTGA, resulting in a 450-bp band for the wild type and a 250-bp band for the targeted AIM2 knockout when PCR analysis was performed on the genomic DNA (Supplementary Fig. 1). All mice were given free access to food and water *ad libitum*. The experimental groups consisted of 20-25 animals in each group.

### Bilateral Common Carotid Artery Stenosis (BCAS) Mouse Model

Animals were anaesthetized with isoflurane and subjected to BCAS, and subsequent chronic hypoperfusion using microcoils specifically designed for the mouse (microcoil specifications: piano wire diameter 0.08mm, internal diameter 0.18mm, coiling pitch 0.5mm, and total length 2.5mm; Sawane Spring Co Ltd, Japan) was described elsewhere^33^. The left and right common carotid arteries (CCAs) were exposed individually, freed from their sheaths, and a microcoil was twined by rotating it around each CCA. The site of surgery was closed, and the mice were observed and taken care of post-surgery until conscious and recovered to freely access food and water ad libitum. To study the disease progression, the experimental animals were divided into eight time-point groups: Sham, 1, 3, 7, 15, 21, 30 and 60-days BCAS. Sham animals were given a skin incision and their CCAs were exposed. All animals were euthanized at their respective end-point after BCAS for subsequent analysis. For biochemical analysis involving Wild Type control and AIM2 KO mice, the animals were divided into three experimental groups: Sham, 15 and 30-days BCAS. However, only the Sham and 30-days timepoints were selected to challenge the cognitive ability of both WT and AIM2 KO mice during behavioral analysis.

### Measurements of Cerebral Blood Flow by Laser Speckle Contrast Imager

High-resolution Laser Speckle Contrast Imager (PSI system, Perimed Inc.) was used to image cerebral blood perfusion and record cerebral blood flow (CBF) before BCAS (baseline), immediately after BCAS surgery and finally at their respective end-points of BCAS. As shown in Supplementary Fig. 2, regions of interest (ROIs) between the bregma and lambda were selected for overall perfusion in the area of two hemispheres. Body temperature was maintained at 37 ± 0.5 °C, and the skull was shaved and exposed by a midline skin incision. The skull was cleaned gently with sterile phosphate buffered saline (PBS) using a cotton applicator. Finally, the image area was kept moist and a non-toxic silicon oil was applied on the skull, which improved imaging. Perfusion images were acquired using the PSI system with a 70mW built-in laser diode for illumination and a 1388 x 1038 pixels CCD camera installed 10cm above the skull (speed 19Hz, and exposure time 6mSec). Acquired images were analyzed for changes in CBF (cerebral perfusion) using a dedicated PIMSoft program (Perimed Inc.).

### Sample Collection and Processing

Following each time points, mice were euthanized by administering a lethal dose of inhaled carbon dioxide (CO2) and the brains harvested. The different brain areas (cerebral cortex, hippocampus and striatum) were immediately separated on ice and were frozen in dry ice for tissue biochemical analysis (n = 7-8 in each experimental group). A separate group of animals were sacrificed for histological analysis. Mice were deeply anaesthetized, and perfusion fixation though the heart was performed with 25mL of chilled 1xPBS (pH 7.4) then followed by 25mL chilled paraformaldehyde (4%). Once the perfusion was completed, the brains were harvested and placed in vials containing 4% paraformaldehyde solution for immersion-fixation overnight at 4°C (n = 5-7 in each experimental group).

### Immunoblot Analysis

Immunoblot analysis was performed as described by us previously^34^. Briefly, brain tissues were homogenized in lysis buffer and then combined with 2 x Laemelli buffer (Bio-Rad Laboratories, Inc., Hercules, CA, USA). Protein samples were then separated on 7.5 to 12.5% v/v sodium dodecyl sulfate (SDS) gels. The SDS-PAGE gels were transferred onto nitrocellulose membranes to probe for proteins. Next, the nitrocellulose membranes were incubated with the following primary antibodies: NLRP1 (Santa Cruz, sc390133), NLRP3 (Adipogen, AG20B0014), NLRC4 (Millipore, 06-1125), AIM2 (Cell Signaling, #13095), IL-1β (Genetex, GTX74034), IL-18 (Biovision, 5180R-100), Total Caspase-1 and Cleaved Caspase-1 (p33 subunit) (Adipogen, AG20B0042), Cleaved Caspase-1 (p20 subunit) (Cell Signaling, #67314), Casapse-8 (Cell Signaling, #4927), Caspase-11 (Cell Signaling, #14340), Total Caspase-3 (Cell Signaling, #9662), Cleaved Caspase-3 (Cell Signaling, #9664), GSDMD (Cell Signaling, #93709), GSDMD-NT (Cell Signaling, #50928), GSDME (Santa Cruz, sc393162), IBA-1 (Abcam, ab5076), GFAP (Cell Signaling, #12389), and β-actin (Sigma-Aldrich, A5441) overnight at 4°C with agitation. Following primary antibody incubation, membranes were washed three times with 1xTBST before incubating with horseradish peroxidase (HRP)-conjugated secondary antibodies (Goat Anti-Rabbit – Cell Signaling Technology, Danvers, MA, USA; Goat Anti-Mouse – Sigma-Aldrich, St. Louis, MO, USA; Goat Anti-Rat – GE Healthcare Life Sciences, Little Chalfont, UK) for 1hr at 24°C with agitation. Following secondary antibody incubation, membranes were washed three times with 1xTBST, each time for 10 min. The substrate for HRP, enhanced chemiluminescence (ECL) (Bio-Rad Laboratories, Inc., Hercules, CA, USA) was applied before the membranes were imaged using ChemiDocXRS+imaging system (Bio-Rad Laboratories, Inc., Hercules, CA, USA). Quantification of proteins was conducted using Image J software (Version 1.46; National Institute of Health, Bethesda, MD, USA), where protein densitometry was expressed relative to the densitometry of the corresponding β-actin.

### Luxol Fast Blue and Cresyl Violet Staining

Mouse brain tissues were fixed in 10% neutral buffered formalin and then processed into paraffin wax blocks. Coronal sections (4μm thick) were obtained via microtome sectioning. Axonal fibre density of Luxol-fast-blue (LFB) staining was performed to detect the severity of white matter (WM) lesions. Briefly, de-waxed rehydrated sections were immersed in the LFB solution (Abcam, UK) at 37°C overnight. Excess staining was removed by 95% ethanol treatment followed by washing with deionized water. Grey and white matter differentiation was initiated with the treatment of 0.05% aqueous lithium carbonate (Abcam, UK) for 20 seconds, followed by 70% ethanol until the nuclei are decolorized. Sections were immersed in Cresyl Violet solution (Abcam, UK) for 5 min and washed in deionized water. The sections were dehydrated in an ethanol gradient (70 – 100%), and finally cleared in xylene and mounted with a mounting agent. The bright field images were taken under 4X and 60X magnification using an Olympus upright Fluorescence Microscope BX53. The WM lesions were evaluated in five brain regions: the optic tract, internal capsule, caudoputamen, corpus callosum (Medial) and corpus callosum (Paramedian). The severity of the WM lesions was graded as normal (grade 0), disarrangement of the nerve fibres (grade 1), the formation of marked vacuoles (grade 2), and the disappearance of myelinated fibres (grade 3). The severity of neuropathology was scored by three blinded examiners and the number of neurons in CA1, CA2 and CA3 were counted as previously described^35^. The general morphology and the neurodegeneration in the hippocampus of BCAS animals were assessed by Cresyl violet staining.

### Immunofluorescence Analysis

Paraffinized brain sections were cut from sham and BCAS animals and immunostained with primary antibodies against Cleaved Caspase-1 (Affinity Biosciences, AF4022), Cleaved Caspase-3 (Cell Signaling, #9661), GSDMD (Santa Cruz, sc393581), MBP (Cell Signaling, #78896), MAP2 (Millipore, MAB3418), GFAP (Cell Signaling, #3670; #12389), IBA-1 (Wako, 016-26721; Cell Signaling, #17198), CD86 (Santa Cruz, sc-28347), CD206 (eBioscience, 12-2069-42), C3 (Santa Cruz, sc-28294), S100S10 (Invitrogen, PA5-95505), OLIG2 (R&D System, AF2418) and PECAM-1 (BD Pharmingen, 553370). Images were captured with an Olympus FluoView FV1000 (Olympus, Japan) laser scanning confocal microscope using 20x/0.70 air objective and 100x/1.45 oil objective, with 488nm Argon ion and 543nm HeNe laser as the excitation source. Single confocal images were converted to 512 x 512 pixel 12-bit TIFF images.

### Behavioral Paradigm and Training of BCAS Animals

#### Visual Acuity Test

To assess visual detection, mice were placed on a platform within a rotating drum covered with black and white vertical stripes at consistent spatial frequency^36, 37^. The automated drum was then rotated in a clockwise direction for 30sec, and subsequently an anti-clockwise direction for 30sec, after a 15sec intertrial interval. Both larger stripes (2°) and smaller stripes (1°) were tested for all animals before conducting other behavioral tests.

#### Open Field Test

Locomotor activity was measured using an open field test^38^. Each subject was placed in the center of the open field apparatus (20×20×40cm; Clever Sys Inc.; VA, USA). Total distance traveled (cm) and average speed (mm/s) were recorded and tracked via TopScan (Clever Sys Inc.; VA, USA). Data was collected for 35 minutes.

#### Morris Water Maze

Morris water maze was performed 3 weeks after surgery (sham or BCAS), as described previously^39^. Mice were trained 4 times a day at 10-min intervals for 5 consecutive days in this fixed platform training. In each trial, mice were given 60secs to find the platform. Activity of the animal in the water maze was video-tracked by EthoVision software (Noldus Information Technology, Leesburg, VA), and the escape latency was recorded and analyzed. Long term spatial memory was tested on the 6^th^ day, whereby the platform was removed, and each animal was given 60secs in the water maze after being released from four different directions. Time spent around the platform, number of visits to the targeted quadrant and time spent in the targeted quadrant on the probe day were tracked and analyzed. The probe day was designed to fall between 32 to 34 days of BCAS surgery.

### Double Stranded DNA Measurement

Double stranded DNA from the serum of BCAS animals were measured as previously described using Quant-iT™ PicoGreen ^®^ dsDNA Reagent and Kits. Briefly, 20μL of plasma sample collected was diluted with 1X TE solution. Subsequently, equal volume of working solution of Quant-iT™ PicoGreen reagent was added. It was then protected from light and incubated for 5 minutes at room temperature. Samples were excited at 485nm and fluorescence intensity was measured at 520nm using a Synergy HT multi-detection microplate reader (BioTek, Winnooski, VT).

### Statistical Analysis

Experimental data were analyzed by GraphPad Prism 5.02 and 8.0 software (GraphPad Software, San Diego, CA, USA). All values are expressed as mean ± standard error of the mean (S.E.M). One-way Analysis of Variance (ANOVA) was used, followed by a Bonferroni post-hoc test to determine differences between groups. A P-value <0.05 was deemed to be statistically significant. For behavioral data, non-parametric Kruskal-Wallis test was used to determine differences between groups. A P-value <0.05 was deemed to be statistically significant.

## Results

### Chronic cerebral hypoperfusion increases inflammasome signalling

We first evaluated the protein expression of inflammasome receptors over 60 days of BCAS in the cerebral cortex, hippocampus and striatum (Supplementary Figs. 3, 4 & 5). NLRP3, AIM2 and NLRC4 were all increased in the cerebral cortex, while only AIM2 receptor expression was increased in the hippocampus compared to sham controls (Supplementary Fig. 3a-d, 4). NLRP1 expression was increased in the striatum while other inflammasome receptors remained unchanged (Supplementary Figs. 5a & b). Expression of total caspase-8 and precursor IL-1β were increased in the cerebral cortex after BCAS, whereas a decrease in precursor IL-18 was evident, and ASC and total caspases-1 and -11 were unchanged (Supplementary Fig. 3e & f). In the hippocampus, expression of total caspases-1, −8 −11 and precursor IL-18 increased, whereas ASC was decreased after BCAS (Supplementary Fig. 3g & h). ASC, total caspase-8 and precursors for IL-1β and IL-18 were also increased in the striatum following BCAS (Supplementary Fig. 5c & d).

To assess canonical inflammasome activation, expression of cleaved caspase-1 subunits (p33, transient active unit; p20, final by-product) and −8 were examined. After BCAS, elevated expression of cleaved caspase-1 (p20) and −8 was detected in the cerebral cortex (Fig. 1a & b) and also hippocampus (Fig. 1c & d). Mature IL-1β and IL-18 – direct downstream markers of inflammasome activation – were also higher in the cerebral cortex and hippocampus after BCAS versus sham controls (Fig. 1a-d). Expression of non-canonical cleaved caspase-11 was also elevated in the cerebral cortex and hippocampus after BCAS (Fig. 1a-d), whereas only cleaved caspase-8, IL-1β and IL-18 were increased in the striatum (Supplementary Fig. 6). Overall, the data reveal that inflammasome activation occurs in the brain in a spatial-temporal manner as a result of chronic cerebral hypoperfusion.

**Figure 1:**
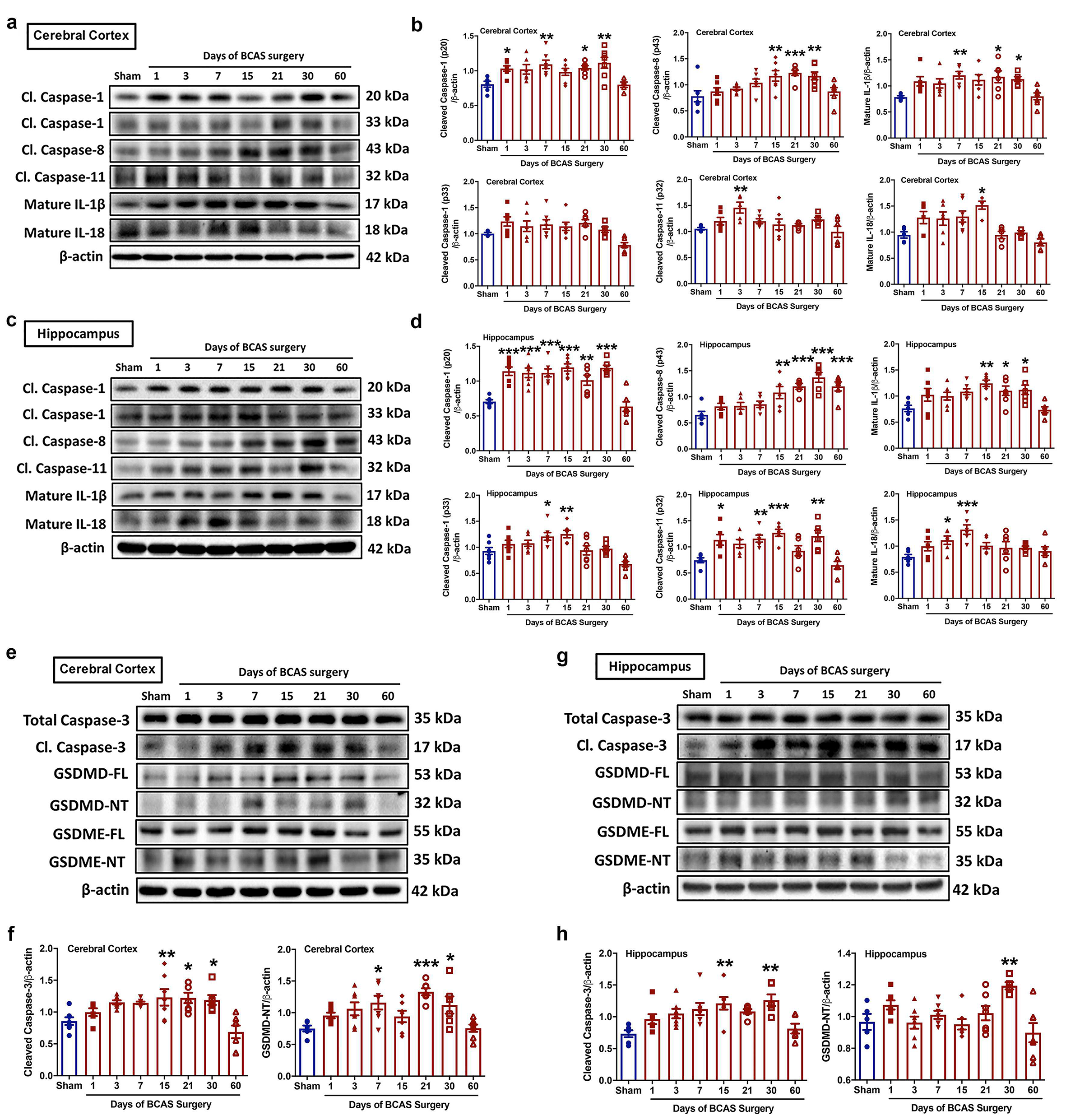
Effect of chronic cerebral hypoperfusion on inflammasome activation and cell death in the cerebral cortex and hippocampus over time following BCAS. (a & b), representative immunoblots and quantification illustrating increased levels of activated inflammasome effector proteins such as cleaved caspase-1 (p20), −8, and −11, and maturation of downstream effector targets, IL-1β and IL-18, in the cerebral cortex. (c & d), representative immunoblots and quantification illustrating increased levels of activated inflammasome effector proteins such as cleaved caspase-1 (p20, p33), −8, and −11, and maturation of downstream effector targets, IL-1β and IL-18, in the hippocampus. (e & f), representative immunoblots and quantification illustrating increases in the expression of apoptotic marker cleaved caspase-3 and pyroptotic marker N-terminal GSDMD in the cerebral cortex. (g & h), representative immunoblots and quantification illustrating increases in the expression of apoptotic marker cleaved caspase-3 and pyroptotic marker N-terminal GSDMD in the hippocampus. β-actin was used as a loading control. Data are represented as mean ± S.E.M. n=6-7 mice in each experimental group. *P<0.05 compared with Sham; **P<0.01 compared with Sham; ***P<0.001 compared with Sham. Abbreviations: BCAS, bilateral common carotid artery stenosis; Cl, cleaved; FL, full length; NT, N-terminal; GSDMD, gasdermin D; GSDME, gasdermin E.

### Chronic cerebral hypoperfusion promotes apoptosis and pyroptosis

We next examined cell death mechanisms associated with inflammasome activation. Thus, we assessed the expression of cleaved caspase-3 and GSDMD-NT, along with their precursor proteins after BCAS (Fig. 1e-h; Supplementary Fig. 7 & 8). Our data showed that in the cerebral cortex (Fig. 1e & f) and hippocampus (Fig. 1g & h) the expression of cleaved caspase-3 and GSDMD-NT were increased following BCAS compared to sham controls. Pyroptotic cell death (i.e. GSDMD-NT expression) was more prominent in the cerebral cortex following BCAS (Fig. 1e & f), whereas only apoptosis (i.e. increased cleaved caspase-3 expression) was detected in the striatum following BCAS (Supplementary Fig. 8).

We also assessed inflammasome-associated secondary pyroptosis/necrosis, which can be induced by cleaved caspase-3, as indicated by expression of N-Terminal Gasdermin E (GSDME-NT). Similar to GSDMD-NT, inflammasome-generated GSDME-NT can permeabilize nuclear and plasma membranes and mitochondria, linking inflammasome activation to cell death^16^. We found GSDME-NT to be increased in the cerebral cortex, but not in the hippocampus or striatum following BCAS (Supplementary Fig. 7 & 8). Expression patterns of inflammasome-related proteins and pathways following BCAS-induced chronic cerebral hypoperfusion are summarized in Supplementary Fig. 9. Overall, the data indicate that BCAS induces differential priming of inflammasome components, inflammasome activation, and induction of apoptosis and pyroptosis pathways in the brain in a spatial-temporal manner.

### Chronic cerebral hypoperfusion induces glial cell activation, white matter lesions and hippocampal neuronal death

We next analyzed for associations between inflammasome activity and hallmark VaD pathologies involving glial cell activation, white matter integrity and neuronal loss. Expression of ionized calcium binding adaptor molecule 1 (Iba1) and glial fibrillary acidic protein (GFAP) indicated activation of microglia and astrocytes, respectively (Fig. 2a-d). Following BCAS, we found increased Iba-1 in the cerebral cortex (Fig. 2a & b) and increased GFAP in the hippocampus (Fig. 2c & d), and both markers increased in the striatum (Supplementary Fig. 10a & b). Subsequently, colocalization of both Iba1 and GFAP with respective polarization markers CD86, CD206, complement component 3 (C3) and S100A10 revealed activation of mainly M1 (CD86 positive) and M2 (CD206 positive) microglia upon chronic cerebral hypoperfusion in the cerebral cortex and hippocampus (Fig. 2e-h; Supplementary Fig. 11a-h) and striatum (Supplementary Fig. 10c-f). Myelin basic protein (MBP) and Luxol fast blue staining revealed myelin integrity, while cresyl violet and microtubule-associated protein 2 (MAP2) staining enabled assessment of neuronal loss in the hippocampus. Staining for MBP and MAP2 was reduced in the cerebral cortex (Fig. 2i & k), hippocampus (Fig. 2j & l) and striatum (Supplementary Fig. 10g & h) following 30 days of BCAS.

**Figure 2:**
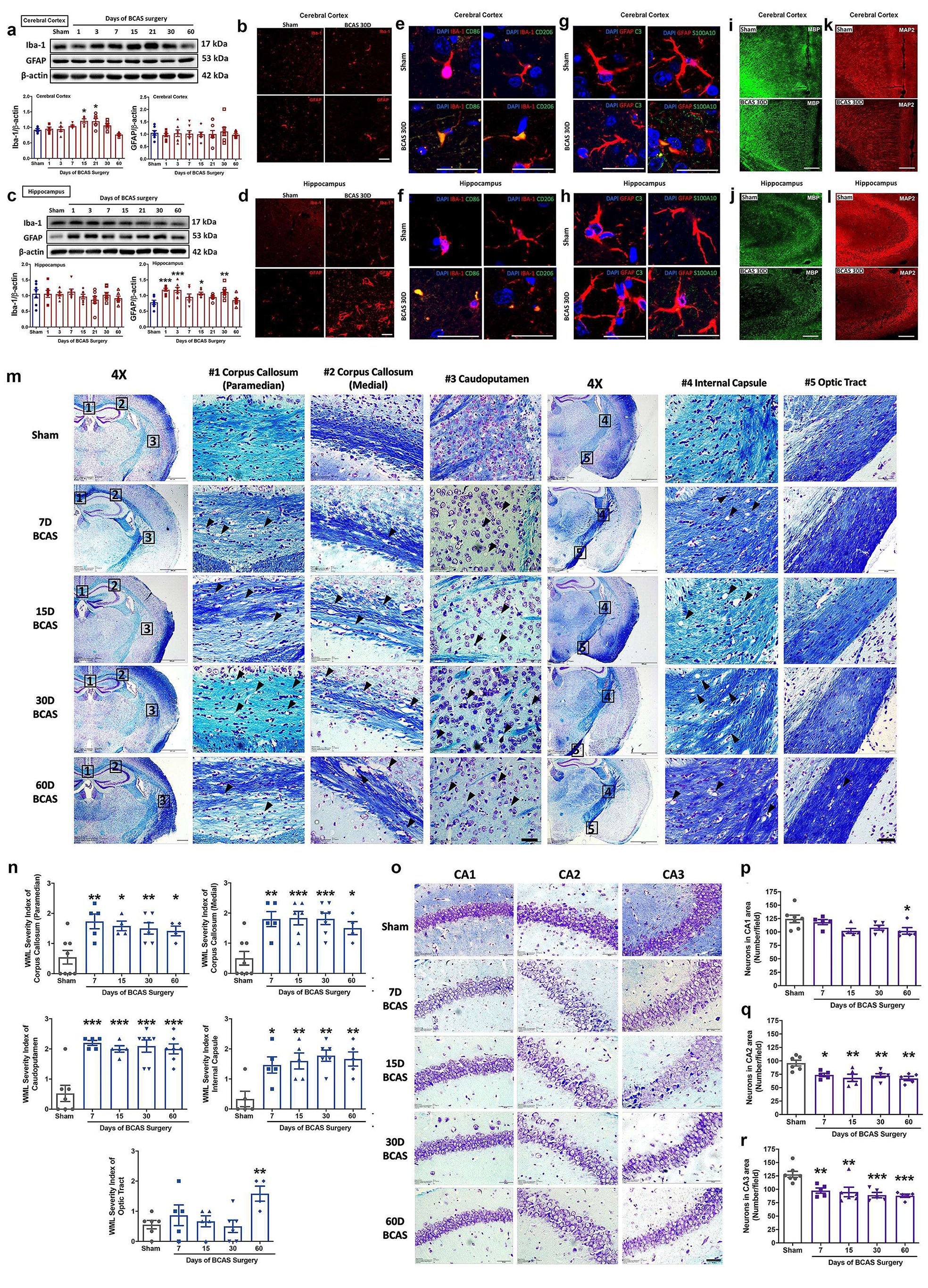
Effect of chronic cerebral hypoperfusion on the levels of glial activation, white matter integrity and hippocampal neuronal density in the cerebral cortex and hippocampus following BCAS. (a), representative immunoblots and quantification illustrating increased microglial activation due to increased levels of Iba-1 in the cerebral cortex over time following BCAS. Data are represented as mean ± S.E.M. n=6-7 mice in each experimental group. β-actin was used as a loading control. *P<0.05 compared with Sham. (b), representative immunofluorescence analysis of Iba-1 and GFAP in the cerebral cortex provide supporting evidence of microglial activation following BCAS. Magnification x 100. Scale bar, 20μm. Images were taken under identical exposures and conditions. (c), representative immunoblots and quantification illustrating increased astroglia activation due to increased levels of GFAP in the hippocampus over time following BCAS. Data are represented as mean ± S.E.M. n=6-7 mice in each experimental group. β-actin was used as a loading control. *P<0.05 compared with Sham; **P<0.01 compared with Sham; ***P<0.001 compared with Sham. (d), representative immunofluorescence analysis of Iba-1 and GFAP in the hippocampus provide supporting evidence of astroglia activation following BCAS. Magnification x 100. Scale bar, 20μm. Images were taken under identical exposures and conditions. (e & f), representative merged immunofluorescence images of DAPI, CD86 and CD206 co-localized within microglia (Iba-1 positive) in the cerebral cortex and hippocampus of WT controls following BCAS. Representative merged immunofluorescence images illustrate the activation of CD86 (M1) and CD206 (M2) positive microglia in the cerebral cortex (e) and hippocampus (f) following BCAS. (g & h), representative merged immunofluorescence images of DAPI, Complement C3 (C3) and S100A10 within astrocytes (GFAP positive) in the cerebral cortex and hippocampus of WT controls following BCAS. Other than colocalization of S100A10 (A2) within astrocytes in cerebral cortex (g), no substantial co-localization of C3 (A1) was observed in the cerebral cortex and hippocampus following BCAS. This illustrates activation of A2 astrocytes in the cerebral cortex (g) upon BCAS. Zoom Magnification x 100. Scale bar, 20μm. (i & j), representative immunofluorescence images illustrating a loss of myelin due to decreased levels of MBP immunoreactivity in the cerebral cortex and hippocampus following BCAS. (k & l), representative immunofluorescence images illustrating a loss of neurons due to decreased levels of MAP2 immunoreactivity in the cerebral cortex and hippocampus following BCAS. Magnification x 20. Scale bar, 120μm. (m & n), representative Luxol fast blue stained images and quantification illustrating disruption of white matter integrity due to increased myelin rarefaction and white matter lesion formation in the corpus callosum (paramedian), corpus callosum (medial), caudoputamen, internal capsule and optic tract over time following BCAS. The severity of white matter disruption was graded accordingly: Grade 0 = no disruption; Grade 1 = disarrangement of nerve fibers; Grade 2 = formation of marked vacuoles; Grade 3 = disappearance of myelinated fibers. Images were taken under identical exposures and conditions. (o-r), representative crystal violet images and quantification illustrating loss of Nissl positively stained neurons in hippocampal CA1, CA2 and CA3 regions over time following BCAS. Data are represented as mean ± S.E.M. n=5-7 mice in each experimental group. *P<0.05 compared with Sham; **P<0.01 compared with Sham; ***P<0.001 compared with Sham. Magnification x 60. Scale bar, 20 μm. Images were taken under identical exposures and conditions. Abbreviations: BCAS, bilateral common carotid artery stenosis; CD86, cluster of differentiation 86; CD206, cluster of differentiation 206; C3, complement component 3; GFAP, glial fibrillary acidic protein; Iba-1, ionized calcium binding adaptor molecule-1; MBP, myelin basic protein; MAP2, microtubule-associated protein 2; S100A10, S100 calcium-binding protein A10.

White matter integrity of five brain regions was assessed after 7-60 days of hypoperfusion, and found to be disrupted in the corpus callosum (Paramedian and Medial), caudoputamen, internal capsule and optic tract (Fig. 2m). All five areas exhibited time-dependent rarefaction of white matter (Fig. 2n) from 7 days, although the white matter lesion (WML) index was not increased in the optic tract until 60 days of BCAS (Fig. 2m & n). BCAS-induced neuronal loss was evident in hippocampal CA1, CA2 and CA3 areas (Fig. 2o-r). Whereas hippocampal sections from sham controls showed normal neuronal cell bodies with distinct nuclei, nucleoli, and densely packed neurons in all three hippocampal areas, there were signs of widespread neuronal loss evident after BCAS (Fig. 2o-r). Degeneration was more pronounced in the hippocampal CA2 and CA3 areas with severe atrophy from 7 days (Fig. 2o-r).

### AIM2 activation mediates apoptosis and pyroptosis during chronic cerebral hypoperfusion

As AIM2 was upregulated in the cerebral cortex and hippocampus, we postulated its involvement in BCAS-induced brain injury. Analysis of serum revealed an increase in cell-free double stranded DNA (dsDNA) levels following BCAS, especially after 15 days (Fig. 3a). As dsDNA is the only known ligand that activates the AIM2 receptor^22, 24^, the finding supports a potential involvement of the AIM2 inflammasome in BCAS-induced brain injury.

**Figure 3:**
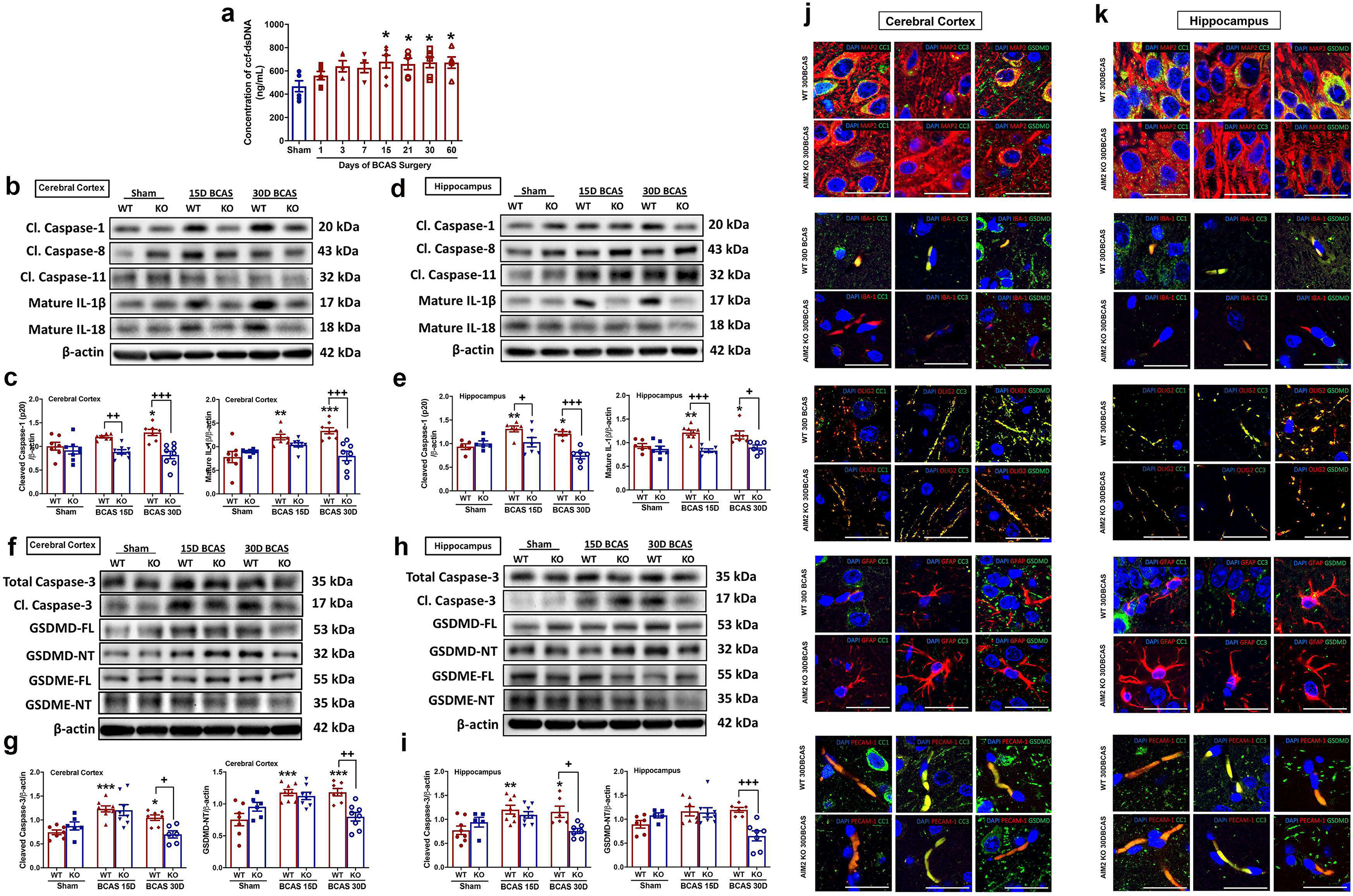
Effect of chronic cerebral hypoperfusion on inflammasome-mediated programmed cell death in AIM2 KO mice following BCAS. (a), quantification illustrating increased serum concentrations of the AIM2 receptor ligand, cell free double stranded DNA (dsDNA), over time in wild-type mice following BCAS. Data are represented as mean ± S.E.M. n=5-7 mice in each experimental group. *P<0.05 compared with Sham. (b-e), representative immunoblots and quantification illustrating suppression of inflammasome activation due to decreased expression of cleaved caspase-1 (p20) at Day 15 and 30, and maturation of IL-1β cytokine production at Day 30 in the cerebral cortex (b & c) and hippocampus (d & e) of AIM2 KO mice compared to WT controls following BCAS. (f-i), representative immunoblots and quantification illustrating attenuation of apoptotic and pyroptotic cell death due to decreased expression of cleaved caspase-3 and GSDMD-NT, respectively, at Day 30 in the cerebral cortex (f & g) and hippocampus (h & i) of AIM2 KO mice compared to WT controls following BCAS. β-actin was used as a loading control. Data are represented as mean ± S.E.M. n=5-8 mice in each experimental group. *P<0.05 compared with WT Sham; **P<0.01 compared with WT Sham; ***P<0.001 compared with WT Sham. ^+^P<0.05 compared with WT BCAS; ^++^P<0.01 compared with WT BCAS; ^+++^P<0.001 compared with WT BCAS. (j & k), representative merged immunofluorescence images of DAPI, cleaved caspase-1 p10 (CC1), cleaved caspase-3 (CC3) and GSDMD co-localized within neurons (MAP2 positive), microglia (Iba-1 positive), oligodendrocytes (OLIG2 positive) and endothelial cells (PECAM-1 positive) in the cerebral cortex and hippocampus of WT controls following BCAS. No substantial co-localization of cleaved caspase-1 p10, cleaved caspase-3 and GSDMD was observed in astrocytes (GFAP positive) in the cerebral cortex and hippocampus of WT controls following BCAS. (j & k), representative merged immunofluorescence images illustrate a reduction in inflammasome activation, and apoptotic and pyroptotic cell death due to decreased expression levels of cleaved caspase-1, cleaved caspase-3 and GSDMD, respectively, in both neurons and microglia in the cerebral cortex (j) and hippocampus (k) in AIM2 KO mice compared to WT controls following BCAS. Zoom Magnification x 100. Scale bar, 20 μm. Images were taken under identical exposures and conditions. Abbreviations: BCAS, bilateral common carotid artery stenosis; Cl, cleaved; FL, full length; NT, N-terminal; WT, wild-type; KO, knock out; CC1, cleaved caspase-1; CC3, cleaved caspase-3; GSDMD, Gasdermin D; MAP2, microtubule-associated protein 2; Iba-1, ionized calcium binding adaptor molecule-1; OLIG2, oligodendrocyte transcription factor 2; GFAP, glial fibrillary acidic protein; PECAM-1, platelet endothelial cell adhesion molecule-1.

To further examine the role of AIM2 inflammasome activation on injury following chronic cerebral hypoperfusion, mice with AIM2 deficiency (AIM2 KO) were studied (Supplementary Fig. 1). We first confirmed via laser speckle contrast imaging that cerebral blood flow was equivalent under basal conditions and reduced following BCAS surgery in wild-type (WT) and AIM2 KO mice (Supplementary Fig. 12). This was similar to previously published data from BCAS in mice^29^. This BCAS-induced reduction in flow was sustained to a similar degree for at least 15 days in both genotypes but was slightly higher in AIM2 KO than WT mice at 30 days (Supplementary Fig. 12a-c).

We found no difference in the basal expression levels of cleaved caspase-1, −8, and −11, and both mature IL-1β and IL-18 (Fig. 3b-e) or in the levels of total caspases −1, −8 and −11, or IL-1β and IL-18 precursors in the cerebral cortex and hippocampus of WT and AIM2 KO mice (Supplementary Fig. 13a-d). However, there was evidence of reduced inflammasome activation following BCAS in AIM2 KO mice when compared to WT control (Fig. 3b-e). Of the three active effector proteins, only canonical cleaved caspase-1 expression was reduced in AIM2 KO mice compared to WT control following BCAS (Fig. 3b-e; Supplementary Fig. 14a & b). There was also reduced levels of mature IL-1β and IL-18 in the cerebral cortex and hippocampus of AIM2 KO mice at 15 and 30 days following BCAS (Fig. 3b-e; Supplementary Fig. 14a & b). These data thus indicated less inflammasome activity occurs after BCAS in the absence of the AIM2 receptor, especially mediated by canonical cleaved caspase-1. Furthermore, protein expression of both cleaved caspase-3 and GSDMD-NT was lower at 30 days after BCAS in the cerebral cortex (Fig. 3f & g) and hippocampus (Fig. 3h & i) of AIM2 KO mice, indicating AIM2 inflammasome mediating apoptosis and pyroptosis under chronic cerebral hypoperfusion.

Immunofluorescence studies indicated that cleaved caspase-1 (CC1) activity was lower in AIM2 KO mice in cortical (Fig. 3j; Supplementary Fig. 15a-e & 16a-e) and hippocampal (Fig. 3k; Supplementary Fig. 15f-j & 16f-j) neurons (MAP2 positive) and microglia (Iba-1 positive) compared to WT controls at 30 days following BCAS. Immunoreactivity against cleaved caspase-1 was similar in cortical and hippocampal oligodendrocytes (OLIG2 positive), astrocytes (GFAP positive) and endothelial cells (PECAM-1 positive) of WT and AIM2 KO mice. Cellular specificity of AIM2 inflammasome-mediated cell death following BCAS was assessed by immunoreactivity against cleaved caspase-3 (CC3) and GSDMD in the cerebral cortex (Fig. 3j; Supplementary Fig. 15a-e; Supplementary Fig.17a-e & 18a-e) and hippocampus (Fig. 3k; Supplementary Fig. 15f-j; Supplementary Fig.17f-j & 18f-j). The data indicate that pro-apoptotic cleaved caspase-3 immunoreactivity was lower in cortical (Fig. 3j; Supplementary Fig. 15a-e & 17a-e) and hippocampal (Fig. 3k; Supplementary Fig. 15f-j & 17f-j) neurons (MAP2 positive) and microglia (Iba-1 positive) of AIM2 KO versus WT mice. Similarly, immunoreactivity of pro-pyroptotic GSDMD was reduced in neurons and microglia in the cortex (Fig. 3j; Supplementary Fig. 15a-e & 18a-e) and hippocampus (Fig. 3k; Supplementary Fig. 15f-j & 18f-j) of AIM2 KO mice. These findings were consistent with more neurons and fewer microglial activation in AIM2 KO than WT mice following chronic cerebral hypoperfusion.

### AIM2 KO mice are resistant to microglial activation, myelin breakdown, hippocampal neuronal loss and cognitive deficits following chronic cerebral hypoperfusion

There was a lower expression of Iba-1 in the cerebral cortex and hippocampus of AIM2 KO than WT mice after BCAS (Fig. 4a-f), especially in CD86 positive inflammatory microglial cells (Fig. 4c and f and Supplementary Fig. 19a-h). These results suggest a role for AIM2 in microglial activation and polarization during chronic cerebral hypoperfusion. We next investigated if AIM2 KO mice are protected against BCAS-induced myelin injury. Although sham-operated WT and AIM2 KO mice displayed no difference in protein expression and immunoreactivity of MBP, at 30 days after BCAS, there was lower expression and weaker immunoreactivity for MBP in the cerebral cortex of WT mice than AIM2 KO mice (Fig. 4g-i). A similar difference in MBP immunoreactivity between WT and AIM2 KO was seen in CA2 and CA3 hippocampal regions (Fig. 4j-l). As myelin integrity was healthier in AIM2 KO mice following BCAS, we assessed whether these mice might be resistant to WML. After BCAS, Luxol fast blue staining in WT mice indicated myelin rarefaction and vacuole formation in several WM regions, but a smaller degree of damage was present in the corpus callosum (paramedian or medial), caudoputamen or internal capsule regions of AIM2 KO mice (Fig. 5a-e). Neuronal loss in hippocampal CA1, CA2 and CA3 regions was also less in AIM2 KO compared to WT mice at 15 and 30 days of cerebral hypoperfusion (Fig. 5f-h). These data confirm a deleterious role of the AIM2 inflammasome in mediating VaD pathologies associated with chronic cerebral hypoperfusion.

**Figure 4:**
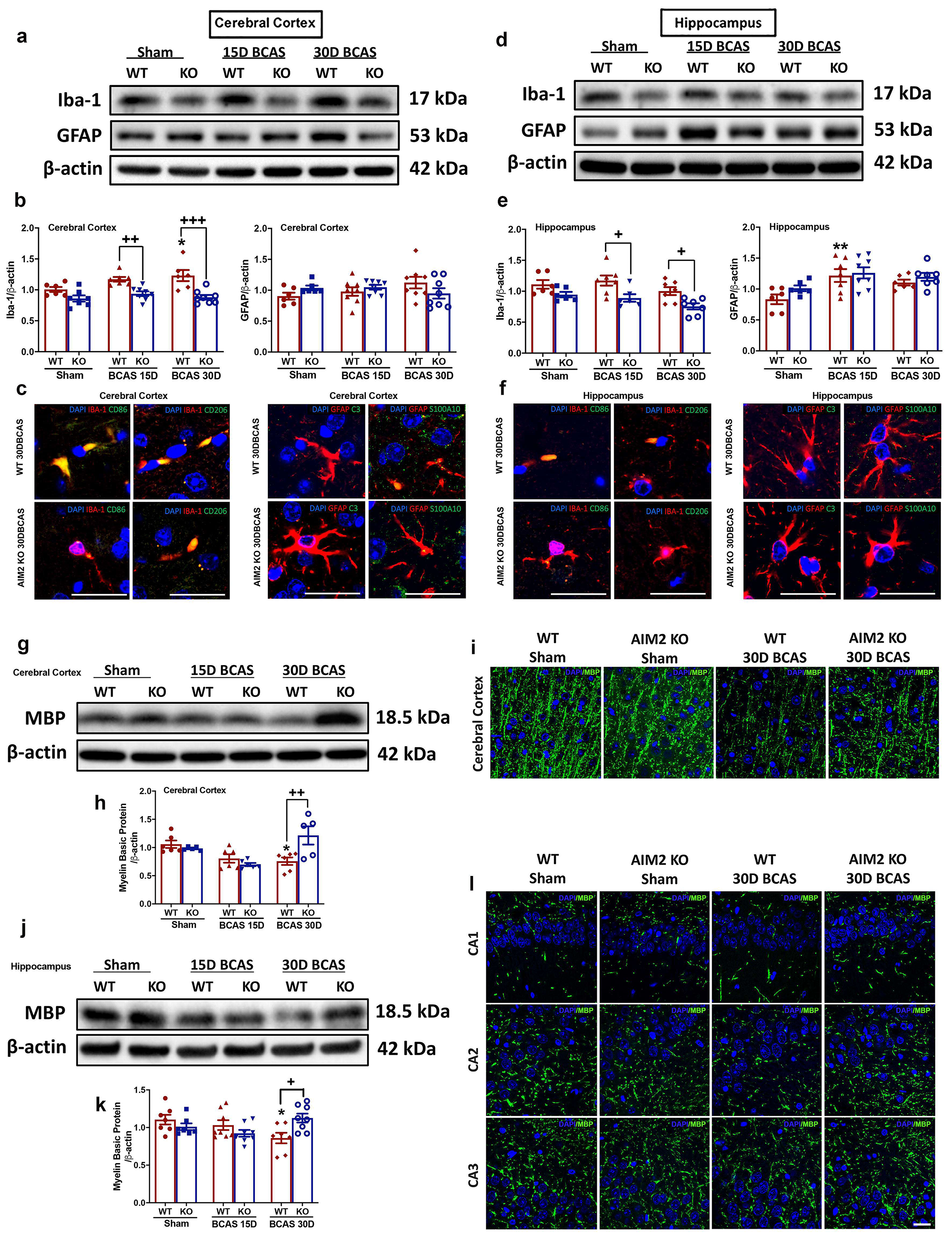
Effect of chronic cerebral hypoperfusion on glial activation and myelin expression in the cerebral cortex and hippocampus of AIM2 KO mice following BCAS. (a & b, d & e), representative immunoblots and quantification illustrating resistance to microglial activation due to decreased expression of Iba-1 in the cerebral cortex and hippocampus of AIM2 KO mice compared to WT controls following BCAS. No effect on astroglia activation was observed as the expression of GFAP in the cerebral cortex and hippocampus remained unchanged in AIM2 KO mice compared to WT controls following BCAS. β-actin was used as a loading control. Data are represented as mean ± S.E.M. n=6-8 mice in each experimental group. *P<0.05 compared with WT Sham; **P<0.01 compared with WT Sham. +P<0.05 compared with WT BCAS; ++P<0.01 compared with WT BCAS; +++P<0.001 compared with WT BCAS. (c & f), representative merged immunofluorescence images of DAPI, CD86 and CD206 co-localized within microglia (Iba-1 positive) in the cerebral cortex and hippocampus of WT controls following BCAS. Representative merged immunofluorescence images illustrate a reduction in M1 (CD86 positive) and M2 (CD206 positive) microglial activation due to decreased expression levels of CD86 and CD206 respectively, in the cerebral cortex (c) and hippocampus (f) in AIM2 KO mice compared to WT controls following BCAS. (c & f), representative merged immunofluorescence images of DAPI, Complement C3 (C3) and S100A10 within astrocytes (GFAP positive) in the cerebral cortex and hippocampus of WT controls following BCAS. Other than colocalization of S100A10 (A2) within astrocytes in the cerebral cortex (c), no substantial co-localization of C3 (A1) was observed in the cerebral cortex (c) and hippocampus (f) of following BCAS. Representative merged immunofluorescence images illustrate a reduction in A2 (S100A10 positive) astrocytes activation due to decreased expression levels of S100A10 in the cerebral cortex (c) in AIM2 KO mice compared to WT controls following BCAS. (g & h, j & k), representative immunoblots and quantification illustrating resistance to myelin loss due to increased expression of MBP in the cerebral cortex (g) and hippocampus (j) in AIM2 KO mice compared to WT controls following BCAS. β-actin was used as a loading control. Data are represented as mean ± S.E.M. n=5-7 mice in each experimental group. *P<0.05 compared with WT Sham; +P<0.05 compared with WT BCAS; ++P<0.01 compared with WT BCAS (i & l), representative immunofluorescence images illustrating resistance to myelin loss due to increased MBP immunoreactivity in the cerebral cortex (i), and hippocampal CA1, CA2 and CA3 regions (l) in AIM2 KO mice compared to WT controls following BCAS. Magnification x 100. Scale bar, 20μm. Images were taken under identical exposures and conditions. Abbreviations: BCAS, bilateral common carotid artery stenosis; CD86, cluster of differentiation 86; CD206, cluster of differentiation 206; C3, complement component 3; GFAP, glial fibrillary acidic protein; Iba-1, ionized calcium binding adaptor molecule-1; KO, knock out; S100A10, S100 calcium-binding protein A10.

**Figure 5:**
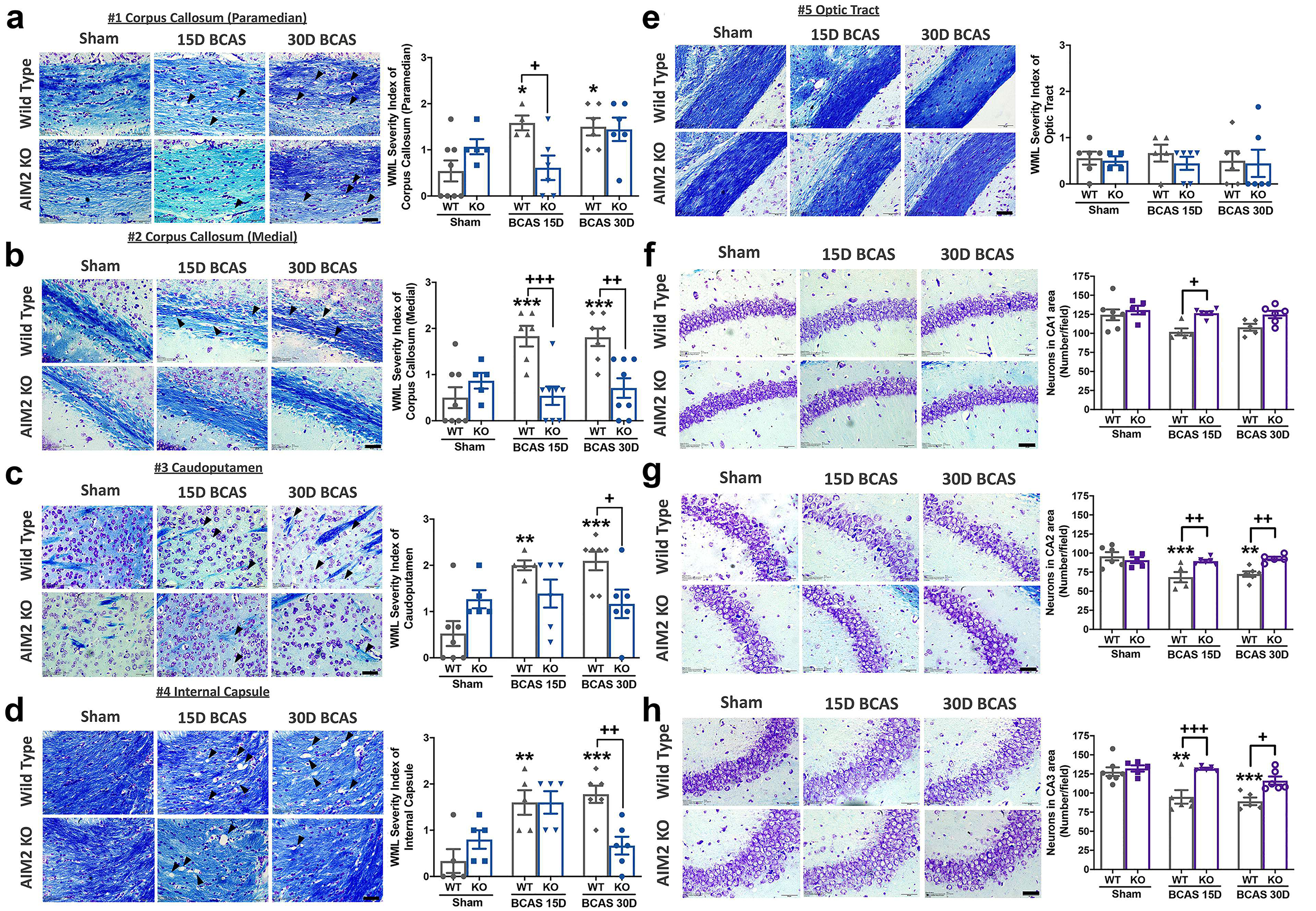
Effect of chronic cerebral hypoperfusion on cortical white matter integrity and hippocampal neuronal density in AIM2 KO mice following BCAS. (a-e), representative Luxol fast blue stained images and quantification illustrating preserved white matter integrity due to decreased myelin rarefaction and white matter lesion formation in the corpus callosum (paramedian), corpus callosum (medial), caudoputamen and internal capsule in AIM2 KO mice compared with WT controls following BCAS. No significant difference in white matter integrity was observed in the optic tract between AIM2 KO mice compared with WT controls following 30 days of BCAS. The severity of white matter disruption was graded accordingly: Grade 0 = no disruption; Grade 1 = disarrangement of nerve fibres; Grade 2 = formation of marked vacuoles; Grade 3 = disappearance of myelinated fibres. (f-h), representative crystal violet images and quantification illustrating increased Nissl positively stained neurons in hippocampal CA1, CA2 and CA3 regions in AIM2 KO mice compared to WT controls following BCAS. Data are represented as mean ± S.E.M. n=5-7 mice in each experimental group. *P<0.05 compared with WT Sham; **P<0.01 compared with WT Sham; ***P<0.001 compared with WT Sham. ^+^P<0.05 compared with WT BCAS; ^++^P<0.01 compared with WT BCAS; ^+++^P<0.001 compared with WT BCAS. Magnification x 60. Scale bar, 20 μm. Images were taken under identical exposures and conditions. Abbreviations: BCAS, bilateral common carotid artery stenosis; WT, wild type; KO, knock out; WML, white matter lesion.

Finally, we tested for a role of the AIM2 inflammasome in BCAS-induced cognitive decline. It was previously shown that BCAS-induced chronic cerebral hypoperfusion in mice resulted in cognitive impairment such as working memory dysfunction and memory loss^29,30^. To examine the effects of AIM2 KO against BCAS-induced cognitive deficits, we conducted the Morris Water Maze and two other control tests (Fig. 6a-h). To ensure that the animals were able to detect the visual cues during the Morris Water Maze, a visual acuity test was conducted. All animals showed a 100% passing rate, and so proceeded to the subsequent tests. In the open field test conducted to investigate explorative and locomotor ability, WT BCAS mice did not have a shorter distance travelled (Fig. 6a) or a lower average speed of movement (Fig. 6b) when compared to WT Sham controls, indicating no impairment of locomotor ability. Similarly, there was also no significant difference in the time spent and the distance travelled in the outer and inner zone among the WT mice, showing no BCAS-induced anxiety effects (Fig. 6c & d). However, it was observed that both sham and BCAS AIM2 KO mice spent more time at the outer zone as compared to the WT controls (Fig. 6c), indicating that AIM2 KO mice generally exhibit higher levels of anxiety independent of BCAS. Several parameters were assessed in the Morris Water Maze performance, including escape latency during five training days. Generally, we observed a shorter time was taken to reach the platform each day across all groups. Similar daily improvement was exhibited in WT or AIM2 KO mice sham controls and AIM2 KO mice following BCAS, whereas WT mice subjected to BCAS took approximately twice as long (Fig. 6e). Following five days of training, spatial memory was assessed whereby WT mice subjected to BCAS spent less time around the platform and significantly fewer visits to the target quadrant than WT controls (Fig. 6f-h). By contrast, AIM2 KO mice subjected to BCAS spent a longer time around the platform and visited the target quadrant more often than WT BCAS mice (Fig. 6f-h). This suggest that AIM2 KO mice exhibit better memory retention as compared to WT controls under similar BCAS conditions. Thus, the behavioral data indicates that BCAS-induced cognitive impairments were reduced in mice deficient in AIM2, suggesting a pivotal role for the AIM2 inflammasome in mediating VaD pathology associated with chronic cerebral hypoperfusion.

**Figure 6:**
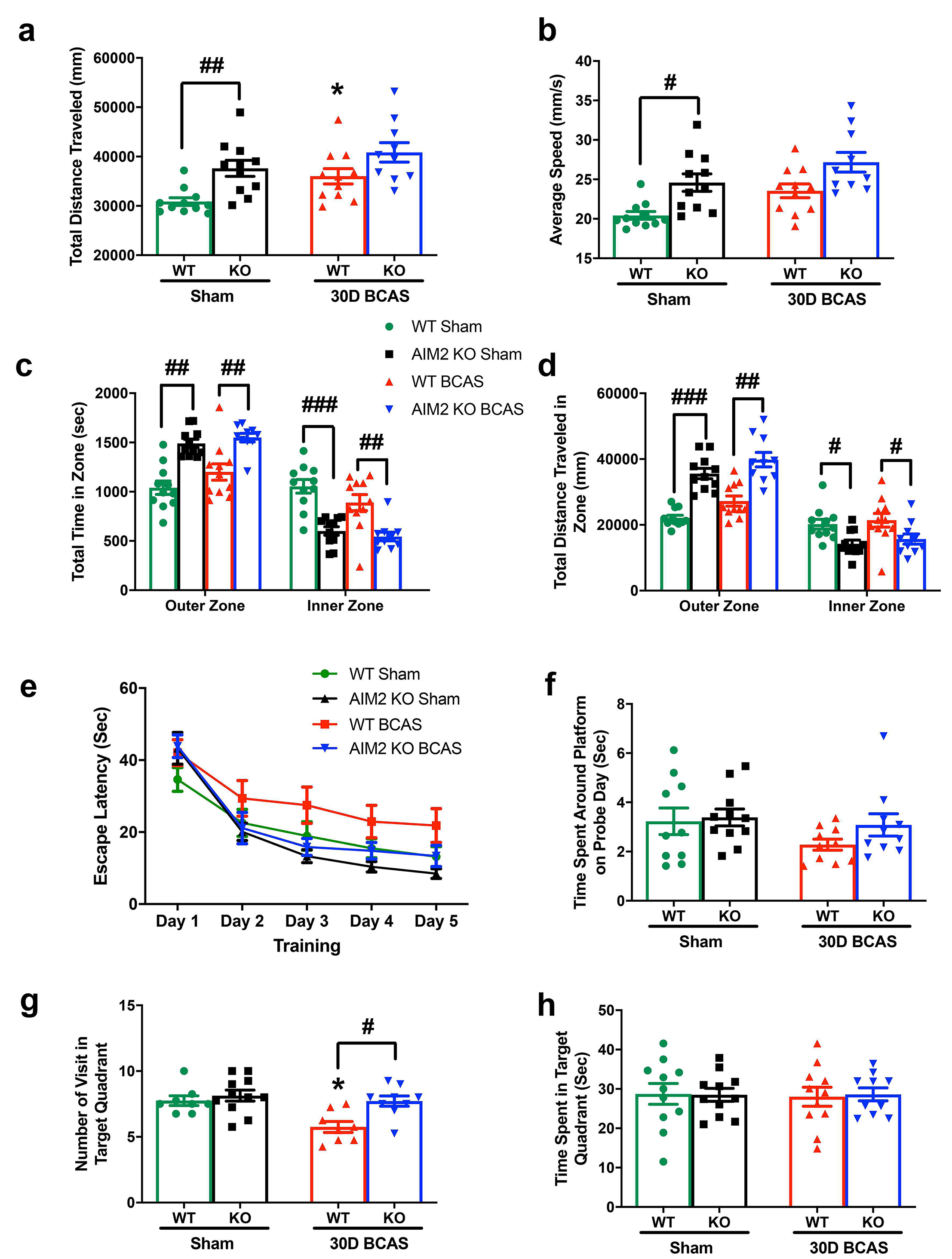
Effect of chronic cerebral hypoperfusion on explorative locomotive behavior and spatial memory in AIM2 KO mice following BCAS. (a-d), quantifications illustrating an open field test conducted on all mice to examine the locomotor ability and level of anxiety by measuring the total distance travelled (mm) and average speed of movement (mm/s); total time (sec) and distance travelled (mm) in the outer and inner zone of the open field. It was shown that WT BCAS mice did not have a shorter distance travelled (mm) (a) and a lower average speed (mm/s) of movement (b) when compared to WT Sham controls, indicating no impairment of locomotor ability. Similarly, there are also no significant difference in the time spent and distance travelled in the outer and inner zone among WT mice, showing no BCAS-induced anxiety effect (c & d). However, it was observed that in both sham and BCAS AIM2 KO mice more time was spent at the outer zone when compared to WT controls (c), indicating that AIM2 KO mice generally exhibit higher levels of anxiety independent of BCAS. (e-h), quantifications illustrating a Morris water maze test conducted on all mice to examine spatial learning and memory by measuring the escape latency (sec), time spent around the platform on probe day (sec), number of visits in the target quadrant and time spent in target quadrant (sec). (e), in general, a reduction in escape latency (sec) was observed in all mice across the training period. Specifically, WT Sham, AIM2 KO Sham and AIM2 KO BCAS groups displayed a trend with a shorter escape latency and steeper declining gradient, which reflected an improved learning ability when compared to the WT BCAS group. (f-h), in general, the AIM2 KO BCAS group displayed a trend with a longer time spent around the platform on probe day and a significantly higher number of visits and time spent in the target quadrant, which reflected retention of spatial memory when compared to the WT BCAS group. Data are represented as mean ± S.E.M. n=9-11 mice in each experimental group. *P<0.05 compared with WT Sham. #P<0.05 compared with WT BCAS; ##P<0.01 compared with WT BCAS; ###P<0.001 compared with WT BCAS. Abbreviations: BCAS, bilateral common carotid artery stenosis; WT, wild type; KO, knock out.

## Discussion

Our findings indicate that inflammasome receptors are expressed in the cortex and hippocampus where the AIM2 inflammasome is activated in response to chronic cerebral hypoperfusion, leading to cellular pathology and cognitive impairment in a mouse model of vascular dementia. Genetic deficiency of AIM2 expression resulted in less brain inflammation, white matter injury, neuronal loss and cognitive decline during cerebral hypoperfusion. These data are the first to indicate that the AIM2 inflammasome promotes neuronal and white matter injury following cerebral hypoperfusion, identifying AIM2 as a key contributor to sterile inflammatory responses in vascular dementia.

Inflammasomes are intracellular multiprotein complexes composed of sensors for various microbial components, viral RNA and damage- or danger-associated molecular patterns (DAMPs) produced during cell injury^40,41^. These innate immune complexes are categorized according to their structural characteristics as either nucleotide-binding domain–like receptors (NLRs) or absent in melanoma 2 (AIM2)-like receptors^41,42^. These receptors oligomerize in response to activation by external stimuli and then recruit the adaptor protein, apoptosis-associated speck-like protein containing a caspase recruitment domain (ASC)^43^. Ultimately, binding of caspase-1 results in its cleavage and activation, and the generation of mature pro-inflammatory cytokines, IL-1β and IL-18, as well as pro-apoptotic cleaved caspase-3, and pro-pyroptotic N-terminal GSDMD and GSDME^44^. While our study established the role of AIM2 in chronic hypoperfusion-induced brain injury, it is still unknown whether other inflammasomes might have a role in WML formation and neuronal cell death following hypoperfusion. Although our data shows increased expression levels of NLRP3 and NLRC4 in brain tissues following BCAS suggesting a potential involvement of other inflammasome complexes, further studies are required to establish the role of these inflammasomes in chronic hypoperfusion-induced brain injury.

We previously demonstrated that inflammasomes contribute to neuronal cell death following ischemic stroke^7^, in part via NF-κB and MAPK signaling^34^. Our present data indicate that chronic cerebral hypoperfusion may also lead to increased expression and activation of NLRP and AIM2 inflammasome receptors in multiple brain areas, especially in the hippocampal region, which is known to be particularly vulnerable to injury following mild ischemia^45^. Cerebral hypoperfusion in the BCAS model used here is known to impact the hippocampus by mechanisms involving ionic imbalance, excitotoxicity, mitochondrial dysfunction and oxidative stress^46–48^. I’m not sure what the purpose of this sentence is.

It is important to elucidate the mechanism(s) by which the AIM2 inflammasome mediates chronic hypoperfusion-induced brain injury. Here we found that the plasma concentration of dsDNA progressively increased following BCAS-induced cerebral hypoperfusion, consistent with the possibility that this ligand promoted AIM2-mediated amplification of brain inflammation and injury. Thus it is plausible that AIM2 functions as a DAMP sensor to promote sterile inflammation in vascular dementia. We found that various white matter regions exhibited injury within 7 days of BCAS, consistent with the known vulnerability of white matter to hypoperfusion. Reduced blood flow to the white matter has been shown to predict the clinical development of WMLs within 18 months^49^. We observed that AIM2 deficient animals subjected to BCAS exhibited greater white matter integrity and lower WML severity scores than WT mice after BCAS. Using magnetic resonance imaging, WMLs appear as white matter hyperintensities (WMH) in VaD patients that arises due to demyelination and axonal loss as well as microglial and endothelial activation^50–52^. In accordance with previous studies, we found that neuronal cell death, loss of myelin basic protein and microglial activation were present in our BCAS mouse model, which were reduced in AIM2 deficient mice. Our data also showed that cerebral blood flow was significantly improved in AIM2 KO mice compared to WT controls following 30 days of BCAS. While we do not know the precise molecular and cellular mechanisms behind this increase, it is possible that reduced inflammasome activation and brain injury may protect the integrity of the neurovasculature to facilitate improved blood flow to the brain.

The sub-regions of the dorsal hippocampus, CA1, CA2 and CA3, are responsible for memory encoding and retrieval^53,54^. Our study showed that neurons in CA2 and CA3 hippocampal regions experienced elevated levels of death sooner in comparison to neurons in the CA1 hippocampal regions during chronic hypoperfusion. Reduced levels of BCAS-induced neuronal loss occurred in hippocampal CA2 and CA3 areas of AIM2-deficient animals when compared to WT controls, which is consistent with reduced inflammasome-mediated cell death in the hippocampal region of AIM2 KO mice, suggesting that the AIM2 inflammasome contributed to neuronal loss in the hippocampus during cerebral hypoperfusion.

The AIM2 receptor is widely expressed in the brain, with its highest levels occurring in microglia under physiological conditions^55^. Furthermore, specifically within the cortex and hippocampus the AIM2 receptor is most abundantly expressed in neurons.^56^. Our data are consistent with a role of AIM2 in these brain resident cells and regions, but we cannot exclude circulating leukocytes as another possible cell mediating the pro-inflammatory actions of AIM2 in the brain in chronic hypoperfusion. Leukocytes contribute to other forms of brain injury and neurodegeneration^57–59^ and their infiltration from the circulation would be expected in association with reduced blood-brain barrier integrity and blood vessel density known to occur in hypoperfusion-induced VaD^60^. However, in this study we have not assessed the role of leukocyte infiltration and related neutrophil extracellular traps (NETs), known as NETosis, as a potential source of AIM2 ligand. It was previously established that NETosis and its associated extracellular dsDNA is known to contribute to the pathogenesis of number of inflammatory diseases^61^. In addition, recent evidences suggest that neutrophils use an inflammasome- and GSDMD-dependent mechanism to activate NETosis, suggesting that further studies are needed to understand the role of NETosis in the pathology of VaD^62^.

Previous studies have recently reported that the AIM2 inflammasome contributes to brain injury and chronic post-stroke cognitive impairment in mice^63^, and to reduced neuroplasticity and spatial memory in a mouse model of AD^20^. Similarly, there is also evidence supporting a role for the NLRP3 inflammasome in cognitive impairment in models of dementia and diabetes^9,64^. Inflammation is known to adversely impact cognitive function, and our data are consistent with inflammasomes promoting an innate immune response to mediate such an effect during cerebral hypoperfusion. Moreover, we observed in our data that AIM2 KO mice generally exhibited higher levels of anxiety independent of BCAS. This was reported by a previous study that established that AIM2 KO mice exhibited increased anxious behaviors and reduced auditory fear memory at the physiological level^65^.

In summary, we have provided substantial evidence that activation of the AIM2 inflammasome plays a key role in promoting brain inflammation, white matter lesions, neuronal cell death and cognitive impairment induced by chronic cerebral hypoperfusion. These effects of AIM2 involve promotion of apoptosis and pyroptosis of cortical and hippocampal neurons. As inflammasome activation plays a major role in a number of inflammatory diseases, there are substantial efforts to develop inflammasome inhibitors. However, application of inflammasome inhibitors is limited in clinical conditions as clinical trials have not yet been completed. While our study establishes that the AIM2 inflammasome may therefore represent a promising therapeutic target for attenuating cognitive impairment in VaD, further research is warranted to develop specific AIM2 inhibitors to test their effect in experimental and clinical studies.

## Acknowledgements

We thank Professor Jenny P. Ting (University of North Carolina, Chapel Hill, Chapel Hill, NC, USA) for providing the AIM2 deficient mice. Supplementary Figure 1 and Supplementary Figure 9a in this article was created using BioRender.

## Funding

This work was supported by the National Medical Research Council Research Grants (NMRC-CBRG-0102/2016 and NMRC/OFIRG/0036/2017), Singapore and the startup fund to TVA from La Trobe University, Melbourne, Australia.

## Author Contributions

Study Conception & Design: T.V.A., L.P., and D.Y.F.; Experiment or Data Collection: T.V.A., L.P., P.W., H.M.L., S.L.F., S.W.K., V.R., S.S., D.Y.F., R.I.V., and N.P.; Data Analysis: L.P., P.W., V.R., S.S., D.Y.F., and T.V.A.; Data Interpretation: L.P., T.V.A., and D.Y.F; Writing-Manuscript Preparation and intellectual input: T.V.A., L.P., D.Y.F., M.B.K., G.R.D., C.G.S., D.C.H., D.G.J., M.K.P.L., C.L.H.C., and L.H.K.L.; Supervision & Administration: T.V.A., D.Y.F., M.K.P.L., and C.L.H.C.

## Conflicts Of Interest

The authors declare that the research was conducted in the absence of any commercial or financial relationships that could be construed as a potential conflict of interest.

## Notes

### Competing Interest Statement

The authors have declared no competing interest.

