## Supplementary Legends for "AIM2 Inflammasome Mediates Hallmark Neuropathological Alterations and Cognitive Impairment in a Mouse Model of Vascular Dementia"

**SUPPLEMENTARY FIGURE LEGENDS**

**S. Figure 1:** **Validation of AIM2 KO mice by PCR analysis.** C57BL/6J mice were used to generate AIM2 KO animals. Genetic modification was done to replace the AIM2 coding region with a neomycin resistance gene. PCR analysis of WT and/or AIM2 KO (with neomycin resistance gene) alleles in the mouse genomic DNA proved that only WT allele (upper band) was present in WT animals and only AIM2 KO allele (lower band) was spotted in KO animals. Both type of alleles were observed in heterozygous AIM2^+/-^ animals.

**S. Figure 2:** **Effect of bilateral common carotid artery stenosis (BCAS) on cerebral blood flow in all experimental WT mice groups.** Laser speckle contrasting imaging was used to monitor the real time cerebral circulation for all experimental groups before, after, and at the terminal timepoint. (a-c), representative contrast images and quantification of basal cerebral blood flow before surgery, effective blood flow reduction after surgery, and the final level of cerebral blood flow at each individual end point demonstrated significant changes in cerebral blood flow following BCAS. Significant blood flow reduction was immediately observed across all experimental groups following BCAS surgery. Although cerebral blood flow levels increased steadily over time (c), a significant degree of cerebral blood flow reduction was maintained till 30 days of BCAS. The rate of blood flow was calculated as an absolute value of cerebral perfusion in perfusion units (PU) using the PeriMed Software. Data are represented as mean ± S.E.M. n = 9-11 mice in each experimental group. *P<0.05 compared with Sham; **P<0.01 compared with Sham; ***P<0.001 compared with Sham.

**S. Figure 3:** **Effect of chronic cerebral hypoperfusion on the expression levels of inflammasome components and both precursor IL-1β and IL-18 in the cerebral cortex and hippocampus over time following BCAS.** (a & b), representative immunoblots and quantification illustrating increases in the expression of NLRP3, AIM2 and NLRC4 inflammasome receptors in the cerebral cortex. (c & d), representative immunoblots and quantification illustrating a decrease in the expression of NLRP3 and increase in the expression of AIM2 inflammasome receptors in the hippocampus. (e & f), representative immunoblots and quantification illustrating increases in the expression of inflammasome components such as total caspase-8 and precursor IL-1β, but a decrease in precursor IL-18 in the cerebral cortex. (g & h), representative immunoblots and quantification illustrating increases in the expression of inflammasome components such as total caspase-1, -8 and -11, and precursor IL-18, while a decrease in the expression of ASC was observed in the hippocampus. β-actin was used as a loading control. Data are represented as mean ± S.E.M. n*=*6-7 mice in each experimental group. *P<0.05 compared with Sham; **P<0.01 compared with Sham; ***P<0.001 compared with Sham. Abbreviations: BCAS, bilateral common carotid artery stenosis.

**S. Figure 4:** **Effect of chronic cerebral hypoperfusion on the expression levels of the NLRP1 inflammasome receptor in the cerebral cortex and hippocampus following BCAS.** Immunoblot quantification illustrating no significant changes in the expression of NLRP1 inflammasome receptors in the cerebral cortex and hippocampus over time following BCAS. β-actin was used as a loading control. Data are represented as mean ± S.E.M. n=6-7 mice in each experimental group. Abbreviations: BCAS, bilateral common carotid artery stenosis.

**S. Figure 5:** **Effect of chronic cerebral hypoperfusion on the expression levels of inflammasome components and both precursor IL-1β and IL-18 in the striatum over time following BCAS.** (a & b), representative immunoblots and quantification illustrating increases in the expression inflammasome receptors such as NLRP1 in the striatum at 15 days of BCAS when compared with Sham. No significant changes in expression were observed for NLRP3, AIM2 and NLRC4 receptors over time. (c & d), representative immunoblots and quantification illustrating increases in the expression of inflammasome components such as ASC, total caspase-1 and -8, and both precursor IL-1β and IL-18 in the striatum over time following BCAS when compared with Sham. β-actin was used as a loading control. Data are represented as mean ± S.E.M. n=5-6 mice in each experimental group. *P<0.05 compared with Sham; **P<0.01 compared with Sham; ***P<0.001 compared with Sham. Abbreviations: BCAS, bilateral common carotid artery stenosis.

**S. Figure 6:** **Effect of chronic cerebral hypoperfusion on inflammasome activation in the striatum over time following BCAS.** Representative immunoblots and quantification illustrating increased levels of activated inflammasome effector protein(s) such as cleaved caspase-8, and maturation of downstream effector targets, IL-1β and IL-18, in the striatum following BCAS. In contrast, decreased levels of activated inflammasome effector proteins such as cleaved caspase-1 p20 and cleaved caspase-11 were observed over time. β-actin was used as a loading control. Data are represented as mean ± S.E.M. n=5-6 mice in each experimental group. *P<0.05 compared with Sham; **P<0.01 compared with Sham; ***P<0.001 compared with Sham. Abbreviations: BCAS, bilateral common carotid artery stenosis; Cl, Cleaved.

**S. Figure 7:** **Effect of chronic cerebral hypoperfusion on the levels of programmed cell death in the cerebral cortex and hippocampus over time following BCAS.** (a), immunoblot quantification illustrating a significant increase in the expression of secondary necrotic marker GSMDE-NT in the cerebral cortex at 21 days of BCAS. In addition, significant increases in the expression of GSDMD-FL was observed in the cerebral cortex at 21 and 30 days of BCAS. (b), immunoblot quantification illustrating a significant decrease in the expression of GSDME-NT in the hippocampus at 30 and 60 days of BCAS. No significant changes were observed for the remaining precursor cell death proteins in the cerebral cortex and hippocampus following BCAS. β-actin was used as a loading control. Data are represented as mean ± S.E.M. n=6-7 mice in each experimental group. *P<0.05 compared with Sham; **P<0.01 compared with Sham; ***P<0.001 compared with Sham. Abbreviations: BCAS, bilateral common carotid artery stenosis; FL, full length; NT, N-terminal; GSDMD, gasdermin D; GSDME, gasdermin E.

**S. Figure 8:** **Effect of chronic cerebral hypoperfusion on the levels of programmed cell death in the striatum over time following BCAS.** (a & b), representative immunoblots and quantification illustrating increases in the expression of apoptotic marker cleaved caspase-3 in the striatum upon 21 days of BCAS. Decrease in pyroptotic marker GSDMD-FL and GSDMD-NT were also observed between 3 and 21 days of BCAS. Moreover, an increase in GSDME-FL was observed at 1 day of BCAS. β-actin was used as a loading control. Data are represented as mean ± S.E.M. n=5-6 mice in each experimental group. *P<0.05 compared with Sham; **P<0.01 compared with Sham; ***P<0.001 compared with Sham. Abbreviations: BCAS, bilateral common carotid artery stenosis; Cl, cleaved; FL, full length; NT, N-terminal; GSDMD, gasdermin D; GSDME, gasdermin E.

**S. Figure 9:** **Schematic diagram of the canonical and non-canonical inflammasome signalling pathway and summary of the levels of proteins involved in inflammasome priming, activation and programmed cell death pathways in the brain following BCAS over time.** (a), the canonical inflammasome pathway is regulated by two signals. The first signal (i.e Priming) involves endogenous extracellular ligands (DAMPs; HMGB1, IL-1α) binding onto pattern recognition receptors (i.e. TLR, RAGE, IFN-γR, IL-1R) on the plasma membrane and activating a number of downstream signalling pathways such as the NF-κB, MAPK, P53 and JAK-STAT pathways to induce gene expression of inflammasome components and both precursor IL-1β and IL-18 that are released into the cytoplasm. The second signal involves the activation of inflammasome receptors that recruits adaptor (i.e. ASC) and effector proteins (i.e. total caspase-1 and -8) to assemble into a multi-protein complex termed an inflammasome. The formation of an inflammasome complex activates and converts total caspase-1 and -8 into active cleaved caspase-1 and -8 via proximity-induced activation. There are three main groups of substrates that are targeted by cleaved caspase-1 and -8. Firstly, cleaved caspase-1 and -8 can cleave both precursor IL-1β and IL-18 into active proinflammatory cytokines, mature IL-1β and IL-18. Secondly, cleaved caspase-1 and -8 can cleave GSDMD-FL into GSDMD-NT that self-oligomerize onto the plasma membrane to form a pore to facilitate the influx of water molecules to induce a lytic form of cell death known as pyroptosis. Thirdly, cleaved caspase-1 and -8 can cleave and activate total caspase-3 into cleave caspase-3 to induce apoptosis. Moreover, cleaved caspase-3 can also initiate another form of cell death known as secondary necrosis by cleaving GSDME-FL into GSDME-NT that similarly self-oligomerize onto the plasma membrane to form a pore to facilitate the influx of water molecules to induce lytic cell death or the mitochondrial membrane to facilitate the leakage of cytochrome c to further exacerbate apoptosis. The non-canonical inflammasome pathway involves total caspase-11 being activated by an endogenous ligand that causes oligomerization and activation of total caspase-11 into cleaved caspase-11 that can directly cleave GSDMD-FL into GSDMD-NT to induce pyroptosis. (b-d), summary chart showing significant changes of protein expression in the inflammasome signalling pathway in the cerebral cortex, hippocampus and striatum over time following BCAS. Most of the significant increases in the cerebral cortex occurred in the latter part of BCAS from 15 to 30 days, while a robust response was observed in the hippocampus throughout the entire duration of BCAS from 1 to 30 days. The striatum demonstrated less of an upregulation of inflammasome components but more of a downregulation of key inflammasome effectors such as cleaved caspase-1 and -11, and pyroptotic marker, GSDMD. Abbreviations: DAMPs, damage associated molecular patterns; HMGB1, high mobility group box protein 1; IL, interleukin; IFN, interferon; TLR, toll-like receptor; RAGE, receptor for advanced glycation end products; NF-κB, nuclear factor kappa-light-chain enhancer of activated B cells; MAPK, mitogen activated protein kinase; JAK/STAT, janus kinase-signal transducer and activator of transcription; Pre, precursor; GSDMD, gasdermin D; GSDME, gasdermin E.

**S. Figure 10:** **Effect of chronic cerebral hypoperfusion on the levels of glial activation, and myelin and neuronal density in the striatum following BCAS.** (a), representative immunoblots and quantification illustrating increased microglial and astroglia activation due to increased levels of Iba-1 and GFAP, respectively, in the striatum over time following BCAS. β-actin was used as a loading control. Data are represented as mean ± S.E.M. n=5-6 mice in each experimental group. *P<0.05 compared with Sham; **P<0.01 compared with Sham; ***P<0.001 compared with Sham. (b), representative immunofluorescence analysis of Iba-1 and GFAP in the striatum provide supporting evidence of microglial and astroglia activation following BCAS. Magnification x 100. Scale bar, 20μm. Images were taken under identical exposures and conditions. (c & d) representative individual and merged immunofluorescence images of DAPI, CD86 and CD206 co-localized within microglia (Iba-1 positive) in the striatum of WT controls following BCAS. Representative merged immunofluorescence images illustrate the activation of M1 (CD86 positive) and M2 (CD206 positive) microglia. (e & f), representative individual and merged immunofluorescence images of DAPI, Complement C3 (C3) and S100A10 within astrocytes (GFAP positive) in the striatum of WT controls following BCAS showing no activation of A1 (C3 positive) and A2 (S100A10 positive) astrocytes. Magnification x 100. Scale bar, 20μm. Images were taken under identical exposures and conditions. (g), representative immunofluorescence images illustrating a loss of myelin due to decreased levels of MBP immunoreactivity in the striatum following BCAS. Magnification x 20. Scale bar, 120μm. Images were taken under identical exposures and conditions. (h), representative immunofluorescence images illustrating a loss of neurons due to decreased levels of MAP2 immunoreactivity in the striatum following BCAS. Magnification x 20. Scale bar, 120μm. Images were taken under identical exposures and conditions. Abbreviations: BCAS, bilateral common carotid artery stenosis; CD86, cluster of differentiation 86; CD206, cluster of differentiation 206; C3, complement component 3; GFAP, glial fibrillary acidic protein; Iba-1, ionized calcium binding adaptor molecule-1; MBP, myelin basic protein; MAP2, microtubule-associated protein 2; S100A10, S100 calcium-binding protein A10.

**S. Figure 11: Effect of chronic cerebral hypoperfusion on glial activation in the cerebral cortex and hippocampus following BCAS.** (a & b, e & f), representative individual and merged immunofluorescence images of DAPI, CD86 and CD206 co-localized within microglia (Iba-1 positive) in the cerebral cortex and hippocampus of WT controls following BCAS. Representative merged immunofluorescence images illustrate the activation of M1 (CD86 positive) and M2 (CD206 positive) microglia in the cerebral cortex (a & b) and hippocampus (e & f) following BCAS. (c & d, g & h), representative individual and merged immunofluorescence images of DAPI, Complement C3 (C3) and S100A10 within astrocytes (GFAP positive) in the cerebral cortex and hippocampus of WT controls following BCAS. Other than colocalization of S100A10 (A2) within astrocytes in the cerebral cortex (d), no substantial co-localization of C3 (A1) was observed in the cerebral cortex (c) and hippocampus (g & h) following BCAS. This illustrates the activation of A2 astrocytes in the cerebral cortex (d) upon BCAS. Magnification x 100. Scale bar, 20μm. Images were taken under identical exposures and conditions. Abbreviations: BCAS, bilateral common carotid artery stenosis; CD86, cluster of differentiation 86; CD206, cluster of differentiation 206; C3, complement component 3; GFAP, glial fibrillary acidic protein; Iba-1, ionized calcium binding adaptor molecule-1; S100A10, S100 calcium-binding protein A10.

**S. Figure 12:** **Effect of chronic cerebral hypoperfusion on cerebral blood flow in AIM2 KO mice over time following BCAS.** Laser speckle contrasting imaging was used to monitor the real time cerebral circulation for WT and AIM2 KO mice before, after, and at the terminal time point. (a-c), representative contrast images and quantification of basal cerebral blood flow before surgery, effective blood flow reduction after surgery, and the final level of cerebral blood flow at the individual end point demonstrated changes in cerebral blood flow following BCAS on WT and AIM2 KO mice. Significant blood flow reduction was immediately observed across all experimental groups following BCAS. Although cerebral blood flow was reduced under 15 and 30 days of BCAS in WT mice, the AIM2 KO mice displayed significantly higher cerebral blood flow levels as compared to WT control at 30 days BCAS. The rate of blood flow was calculated as an absolute value of cerebral perfusion in perfusion unit (PU) using the PeriMed Software. Data are represented as mean ± S.E.M. n = 11-15 mice in each experimental group. ***P<0.001 compared with WT Sham; ^##^P<0.01 compared with KO Sham; ^###^P<0.001 compared with KO Sham; ^+^P<0.01 compared with WT BCAS.

**S. Figure 13:** **Effect of chronic cerebral hypoperfusion on the expression levels of inflammasome components and both precursor IL-1β and IL-18 in the cerebral cortex and hippocampus in AIM2 KO mice following BCAS.** Representative immunoblots and quantification illustrating no significant changes in the expression levels of inflammasome components and both precursor IL-1β and IL-18 in the cerebral cortex and hippocampus of AIM2 KO mice following BCAS. β-actin was used as a loading control. Data are represented as mean ± S.E.M. n=5-8 mice in each experimental group. *P<0.05 compared with Sham; **P<0.01 compared with Sham. Abbreviations: BCAS, bilateral common carotid artery stenosis.

**S. Figure 14:** **Effect of chronic cerebral hypoperfusion on inflammasome activation and programmed cell death in the cerebral cortex and hippocampus of AIM2 KO mice following BCAS.** (a & b), immunoblot quantification illustrating decreased expression of mature IL-18 in the cerebral cortex and hippocampus in AIM2 KO mice compared to WT controls following BCAS. No significant difference were observed in the expression of cleaved caspase-8 and -11 effector proteins between AIM2 KO mice and WT controls following BCAS. (c & d), immunoblot quantification illustrating decreased expression of pyroptotic GSDMD-FL and secondary necrotic GSDME-NT in the cerebral cortex of AIM2 KO mice when compared to WT controls following BCAS. In addition, a decreased expression of apoptotic total caspase-3 was observed in the hippocampus of AIM2 KO mice when compared to WT controls following BCAS. β-actin was used as a loading control. Data are represented as mean ± SEM. n=5-8 mice in each experimental group. *P<0.05 compared with Sham; **P<0.01 compared with Sham; ***P<0.001 compared with Sham. ^+^P<0.05 compared with WT BCAS; ^++^P<0.01 compared with WT BCAS.

**S. Figure 15:** **Effect of chronic cerebral hypoperfusion on the cellular specificity of inflammasome-mediated programmed cell death in the cerebral cortex and hippocampus of AIM2 KO mice following BCAS.** (a-j), representative merged immunofluorescence images of DAPI, cleaved caspase-1 p10, cleaved caspase-3 and GSDMD co-localized within neurons (MAP2 positive), microglia (Iba-1 positive), oligodendrocytes (OLIG2 positive) and endothelial cells (PECAM-1 positive) in the cerebral cortex and hippocampus of WT controls following BCAS. No substantial co-localization of cleaved caspase-1 p10, cleaved caspase-3 and GSDMD was observed in astrocytes (GFAP positive) in the cerebral cortex and hippocampus of WT controls following BCAS. (a & b), representative immunofluorescence images illustrate a reduction in inflammasome activation, and apoptotic and pyroptotic cell death due to decreased expression levels of cleaved caspase-1, cleaved caspase-3 and GSDMD, respectively, in both neurons and microglia in the cerebral cortex in AIM2 KO mice compared to WT controls following BCAS. (f & g), representative immunofluorescence images illustrate a reduction in inflammasome activation, and apoptotic and pyroptotic cell death due to decreased expression levels of cleaved caspase-1, cleaved caspase-3 and GSDMD, respectively, in both neurons and microglia in the hippocampus in AIM2 KO mice compared to WT controls following BCAS. The remaining cell types showed similar cleaved caspase-1, cleaved caspase-3 and GSDMD immunoreactivity between WT controls and AIM2 KO mice following BCAS. Magnification x 100. Scale bar, 20 μm. Images were taken under identical exposures and conditions. Abbreviations: BCAS, bilateral common carotid artery stenosis; WT, wild-type; KO, knock out; CC1, cleaved caspase-1; CC3, cleaved caspase-3; GSDMD, gasdermin D; MAP2, microtubule-associated protein 2; Iba-1, ionized calcium binding adaptor molecule-1; OLIG2, oligodendrocyte transcription factor 2; GFAP, glial fibrillary acidic protein; PECAM-1, platelet endothelial cell adhesion molecule-1.

**S. Figure 16:** **Effect of chronic cerebral hypoperfusion on inflammasome activation in multiple cell types in the cerebral cortex and hippocampus of WT and AIM2 KO mice following BCAS.** (a-j), representative individual immunofluorescence images of DAPI (nucleus marker), cleaved caspase-1 p10 (inflammasome activation marker), and neuronal (MAP2 positive), microglial (Iba-1 positive), oligodendrocyte (OLIG2 positive), astroglia (GFAP positive) and endothelial cell (PECAM-1 positive) immunoreactivity in the cerebral cortex and hippocampus of WT controls and AIM2 KO mice following BCAS. Magnification x 100. Scale bar, 20 μm. Images were taken under identical exposures and conditions. Abbreviations: BCAS, bilateral common carotid artery stenosis; WT, wild-type; KO, knock out; CC1, cleaved caspase-1; CC3, cleaved caspase-3; GSDMD, gasdermin D; MAP2, microtubule-associated protein 2; Iba-1, ionized calcium binding adaptor molecule-1; OLIG2, oligodendrocyte transcription factor 2; GFAP, glial fibrillary acidic protein; PECAM-1, platelet endothelial cell adhesion molecule-1.

**S. Figure 17:** **Effect of chronic cerebral hypoperfusion on apoptotic cell death in multiple cell types in the cerebral cortex and hippocampus of WT and AIM2 KO mice following BCAS.** (a-j), representative individual immunofluorescence images of DAPI (nucleus marker), cleaved caspase-3 (apoptosis marker), and neuronal (MAP2 positive), microglial (Iba-1 positive), oligodendrocyte (OLIG2 positive), astroglia (GFAP positive) and endothelial cell (PECAM-1 positive) immunoreactivity in the cerebral cortex and hippocampus of WT controls and AIM2 KO mice following BCAS. Magnification x 100. Scale bar, 20 μm. Images were taken under identical exposures and conditions. Abbreviations: BCAS, bilateral common carotid artery stenosis; WT, wild-type; KO, knock out; CC1, cleaved caspase-1; CC3, cleaved caspase-3; GSDMD, gasdermin D; MAP2, microtubule-associated protein 2; Iba-1, ionized calcium binding adaptor molecule-1; OLIG2, oligodendrocyte transcription factor 2; GFAP, glial fibrillary acidic protein; PECAM-1, platelet endothelial cell adhesion molecule-1.

**S. Figure 18:** **Effect of chronic cerebral hypoperfusion on pyroptotic cell death in multiple cell types in the cerebral cortex and hippocampus of WT and AIM2 KO mice following BCAS.** (a-j), representative individual immunofluorescence images of DAPI (nucleus marker), GSDMD (pyroptosis marker), and neuronal (MAP2 positive), microglial (Iba-1 positive), oligodendrocyte (OLIG2 positive), astroglia (GFAP positive) and endothelial cell (PECAM-1 positive) immunoreactivity in the cerebral cortex and hippocampus of WT controls and AIM2 KO mice following BCAS. Magnification x 100. Scale bar, 20 μm. Images were taken under identical exposures and conditions. Abbreviations: BCAS, bilateral common carotid artery stenosis; WT, wild-type; KO, knock out; CC1, cleaved caspase-1; CC3, cleaved caspase-3; GSDMD, gasdermin D; MAP2, microtubule-associated protein 2; Iba-1, ionized calcium binding adaptor molecule-1; OLIG2, oligodendrocyte transcription factor 2; GFAP, glial fibrillary acidic protein; PECAM-1, platelet endothelial cell adhesion molecule-1.

**S. Figure 19: Effect of chronic cerebral hypoperfusion on glial activation in the cerebral cortex and hippocampus in AIM2 KO mice following BCAS.** (a & b, e & f), representative individual and merged immunofluorescence images of DAPI, CD86 and CD206 co-localized within microglia (Iba-1 positive) in the cerebral cortex and hippocampus of WT controls following BCAS. Representative merged immunofluorescence images illustrate a reduction in M1 (CD86 positive) and M2 (CD206 positive) microglial activation due to decreased expression levels of CD86 and CD206; respectively, in the cerebral cortex (a & b) and hippocampus (e & f) in AIM2 KO mice compared to WT controls following BCAS. (c & d, g & h), representative individual and merged immunofluorescence images of DAPI, Complement C3 (C3) and S100A10 within astrocytes (GFAP positive) in the cerebral cortex and hippocampus of WT controls following BCAS. Other than colocalization of S100A10 (A2 marker) within astrocytes in the cerebral cortex (d), no substantial co-localization of C3 (A1 marker) was observed in the cerebral cortex (c) and hippocampus (g & h) following BCAS. Representative merged immunofluorescence images illustrate a reduction in A2 astrocyte activation due to decreased expression levels of S100A10 in the cerebral cortex (c) in AIM2 KO mice compared to WT controls following BCAS. Magnification x 100. Scale bar, 20 μm. Images were taken under identical exposures and conditions. Abbreviations: BCAS, bilateral common carotid artery stenosis; CD86, cluster of differentiation 86; CD206, cluster of differentiation 206; C3, complement component 3; GFAP, glial fibrillary acidic protein; Iba-1, ionized calcium binding adaptor molecule-1; S100A10, S100 calcium-binding protein A10.
