## Supplementary figures and images for "AIM2 Inflammasome Mediates Hallmark Neuropathological Alterations and Cognitive Impairment in a Mouse Model of Vascular Dementia"

### S.Figure 1

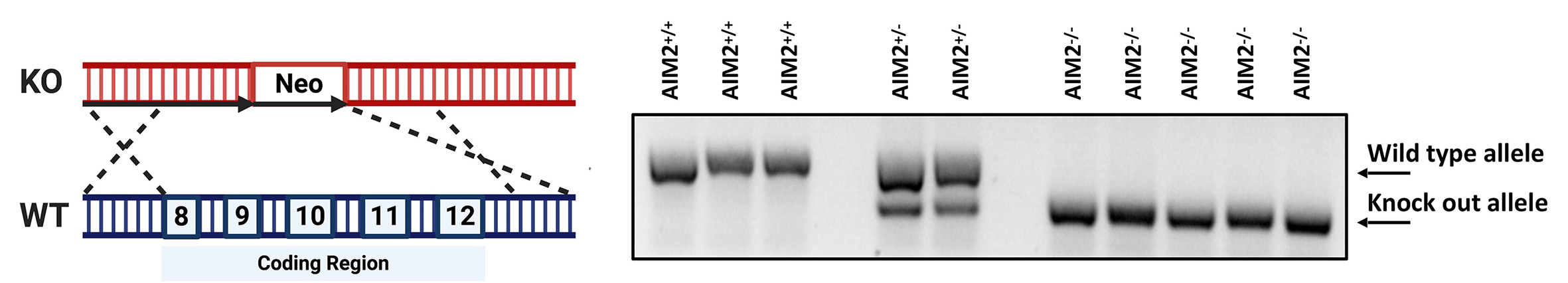

### S.Figure 2

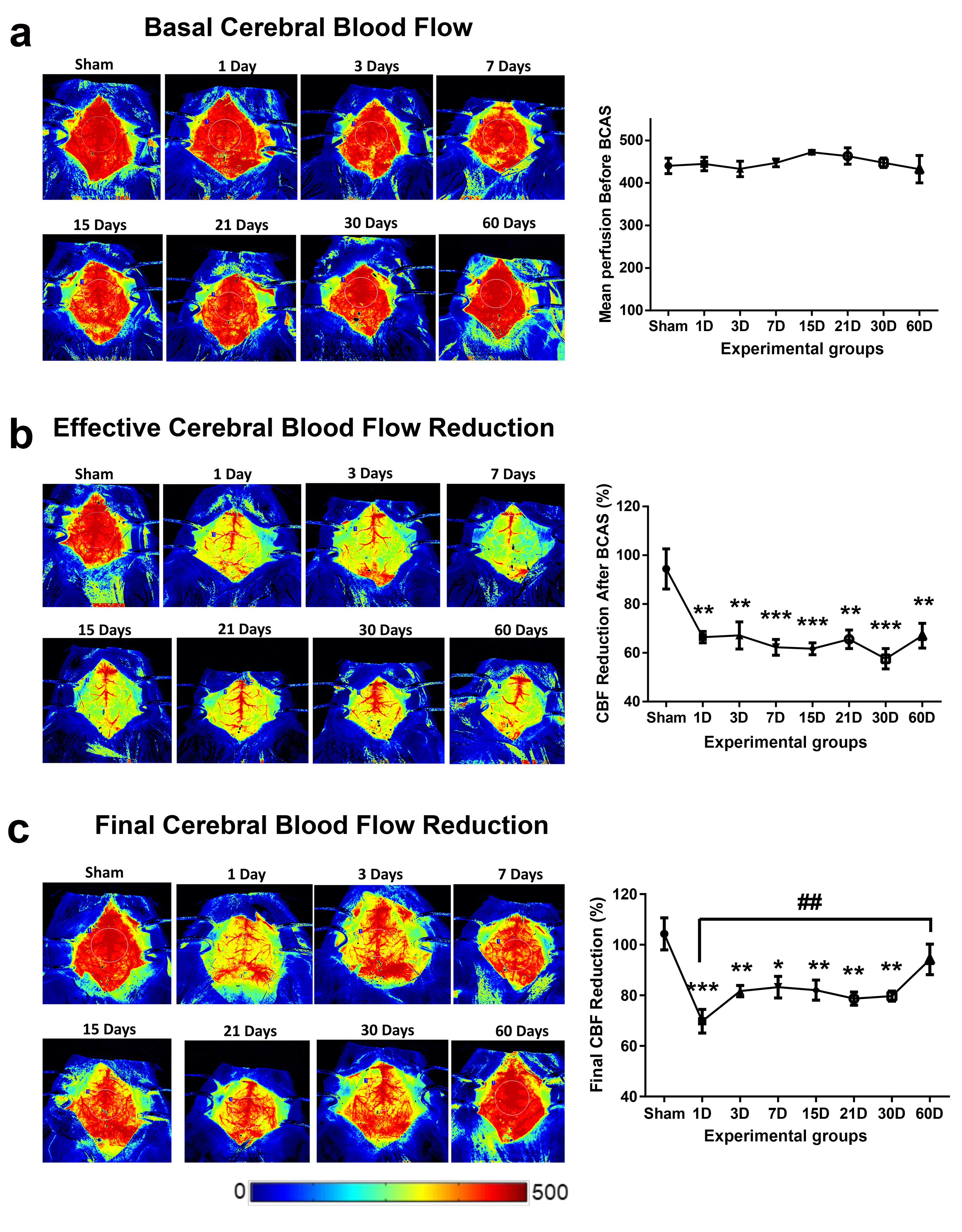

### S.Figure 3

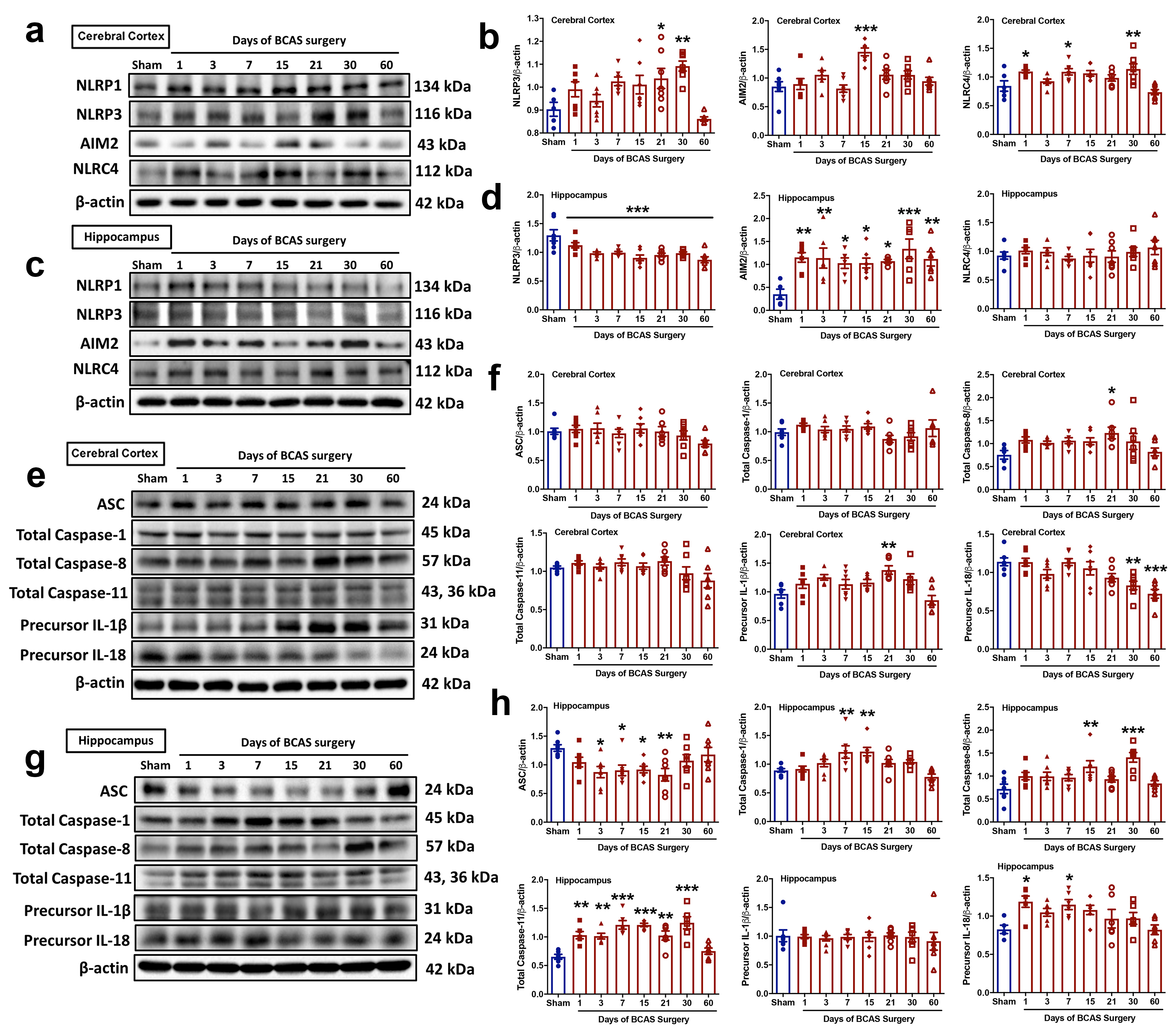

### S.Figure 4

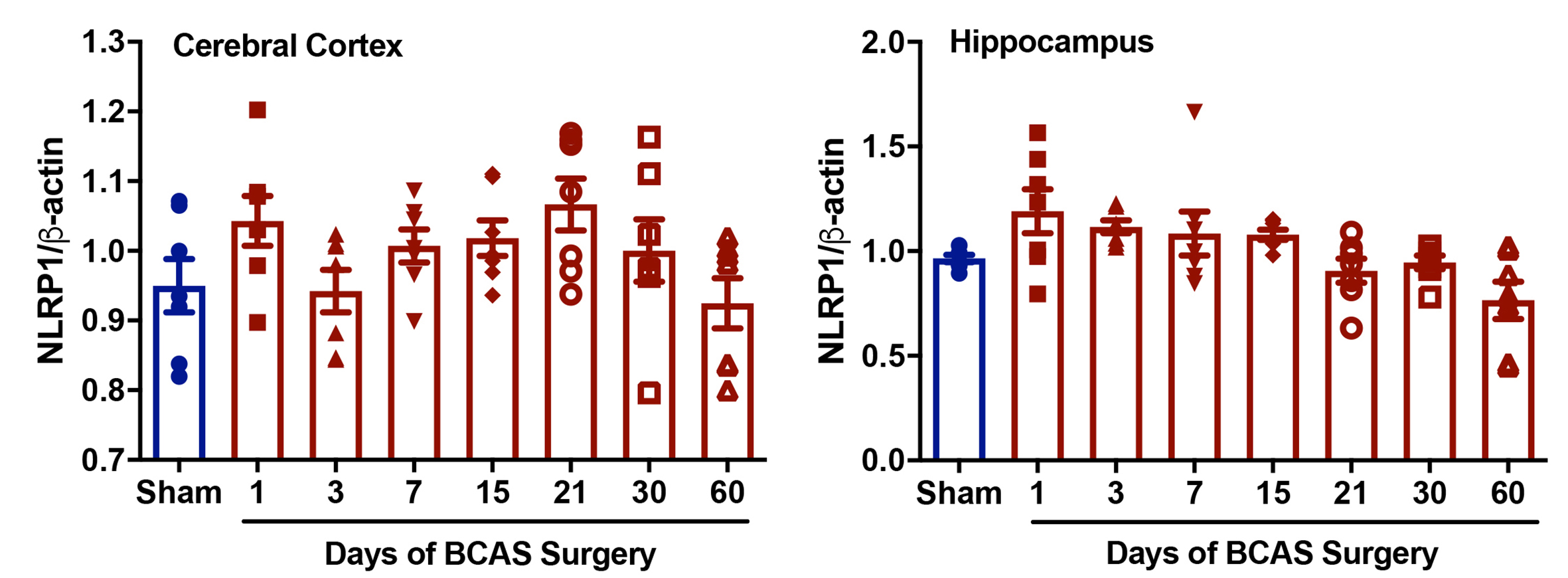

### S.Figure 5

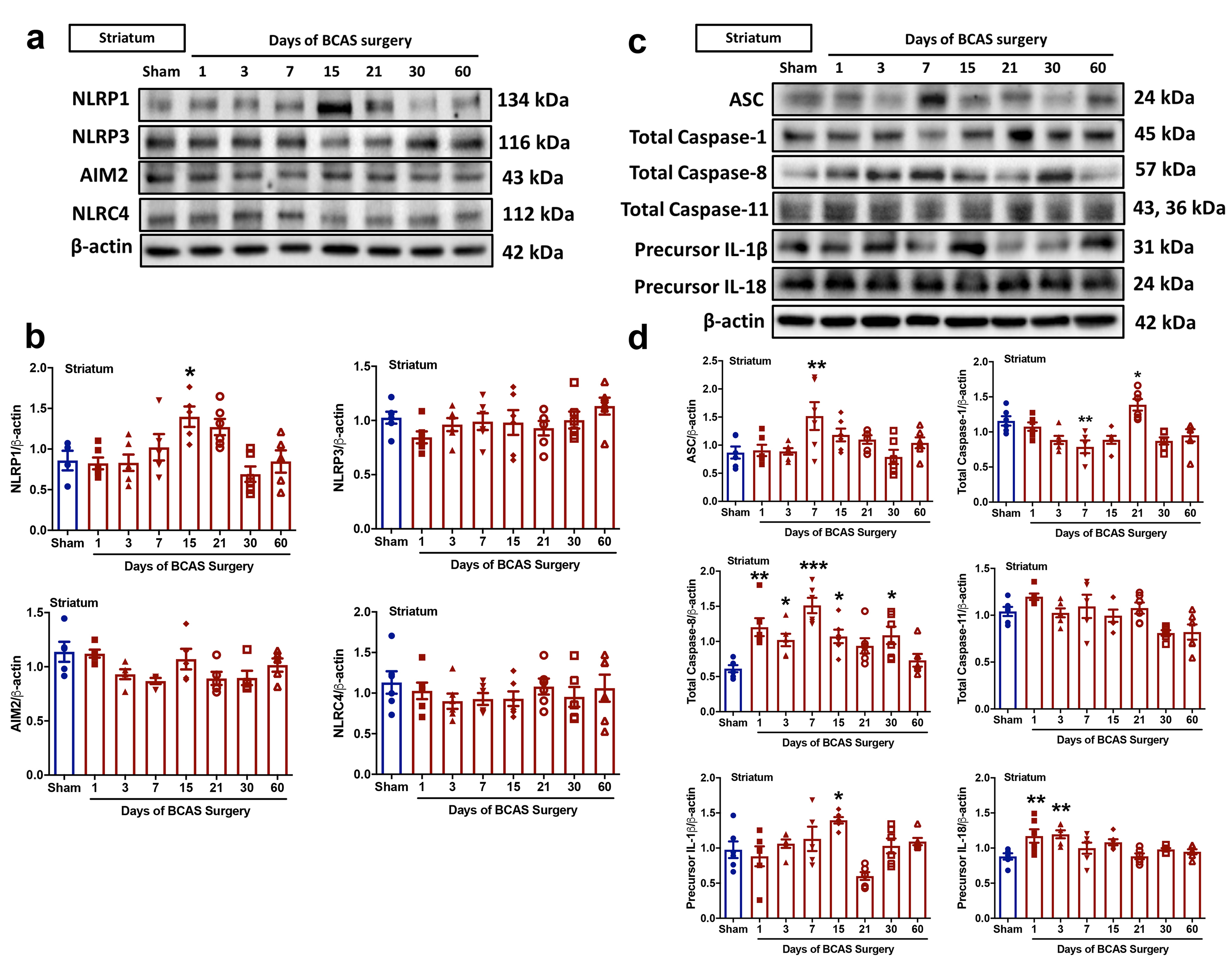

### S.Figure 6

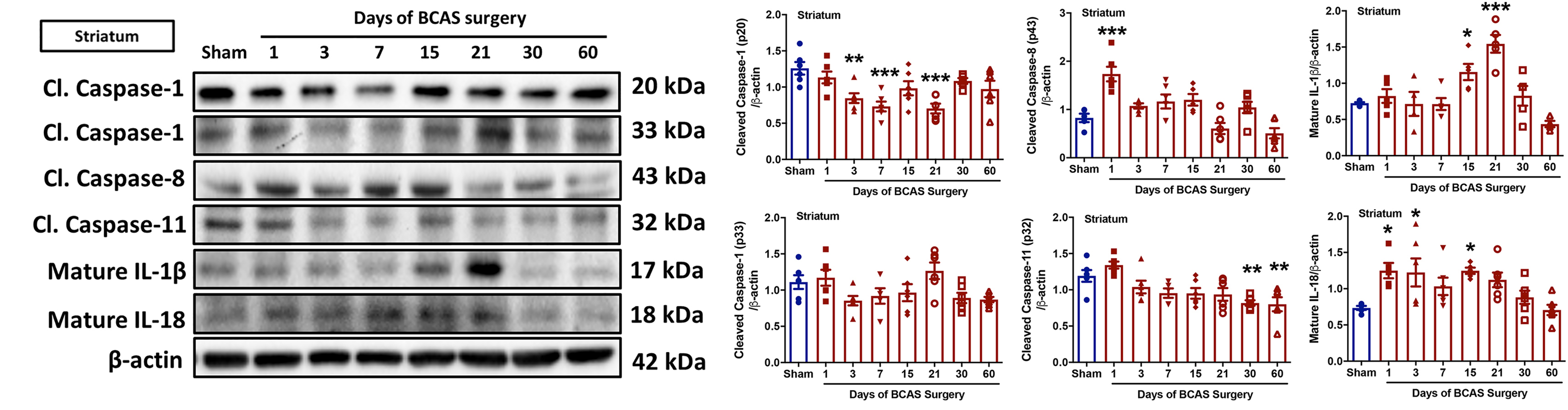

### S.Figure 7

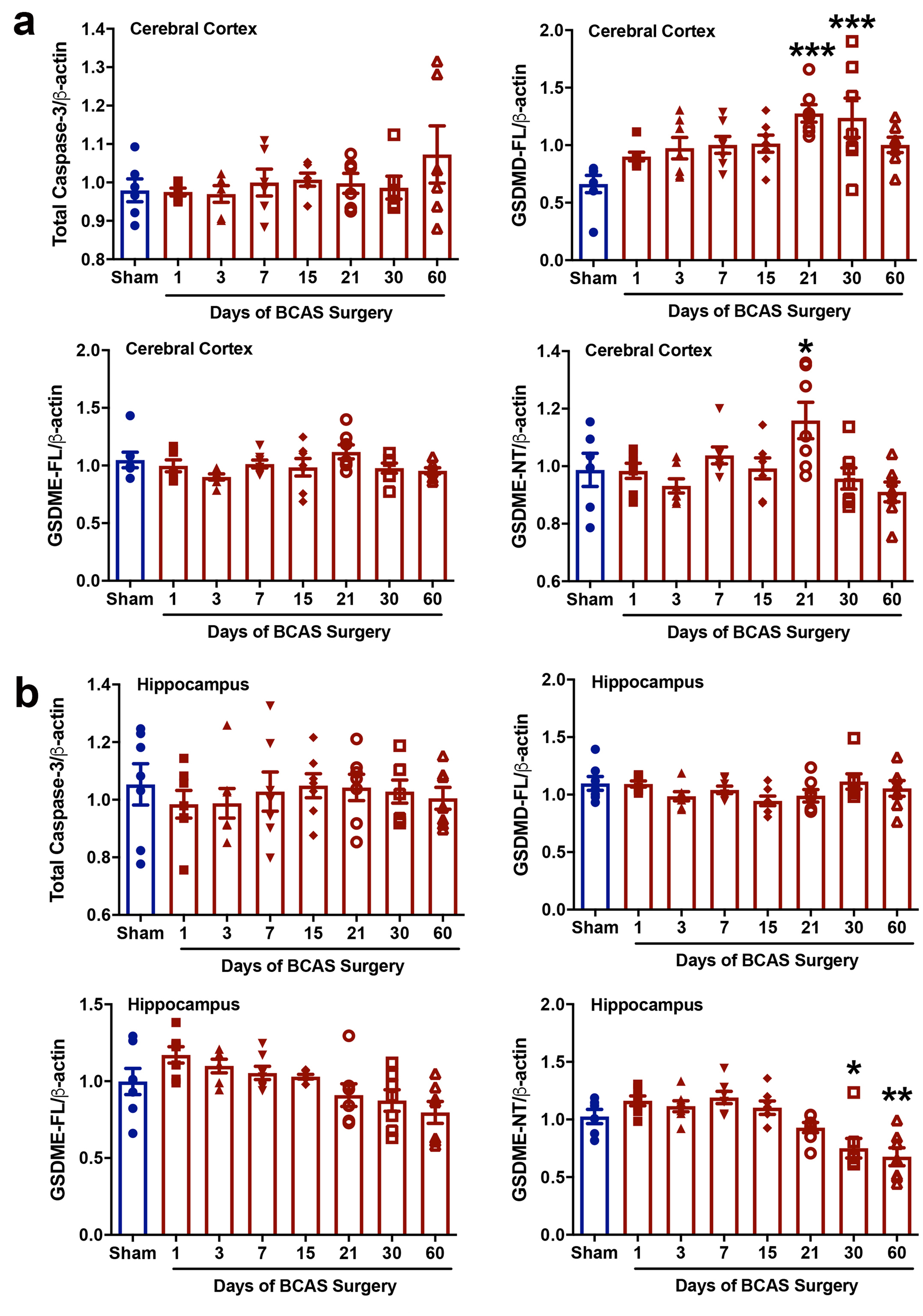

### S.Figure 8

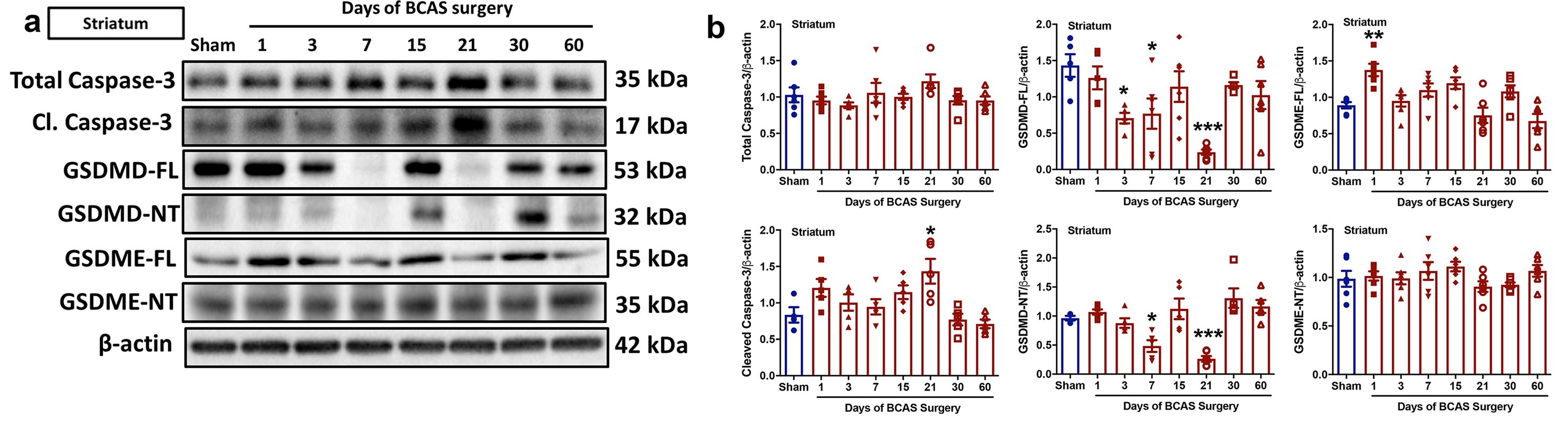

### S.Figure 9

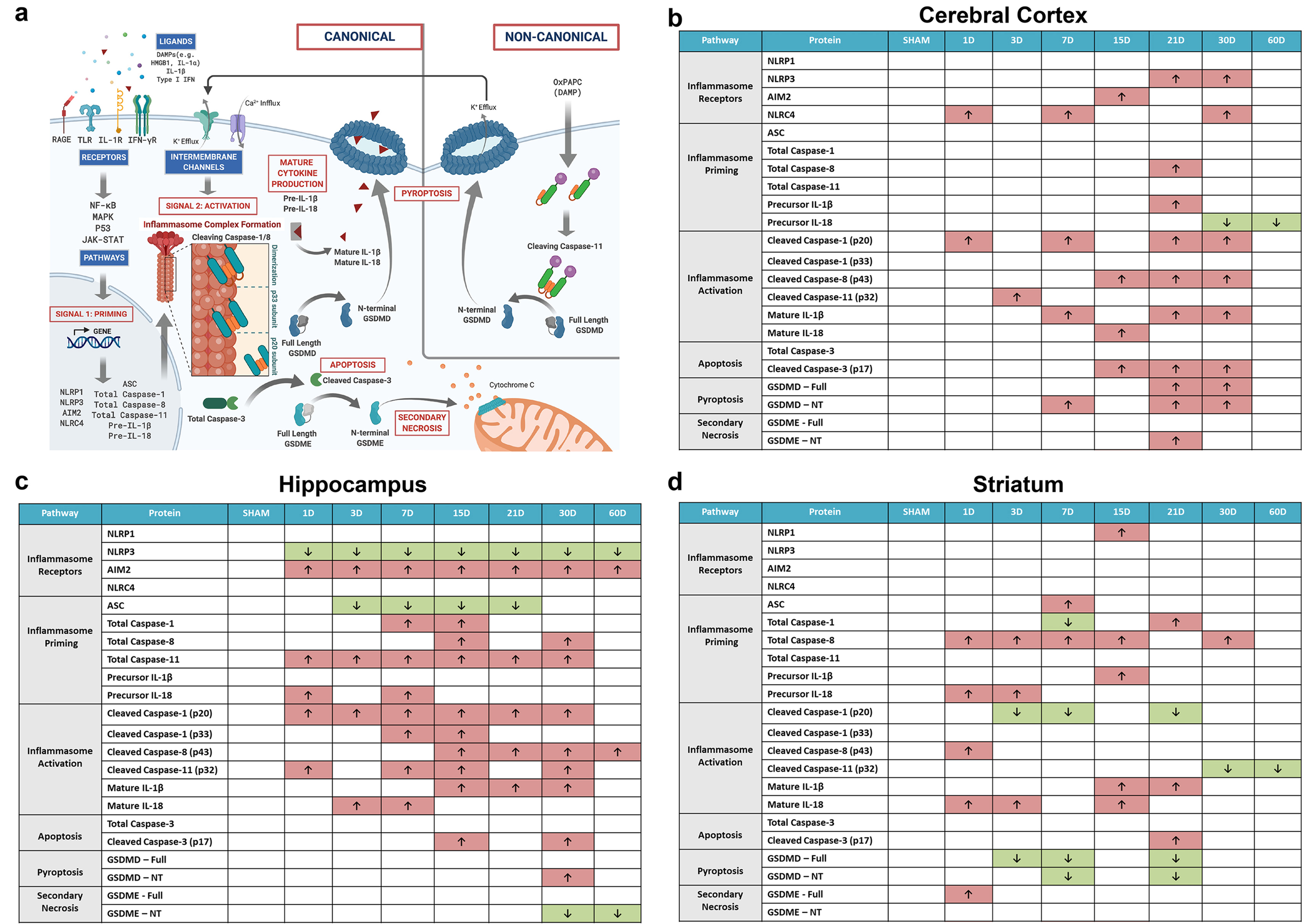

### S.Figure 10

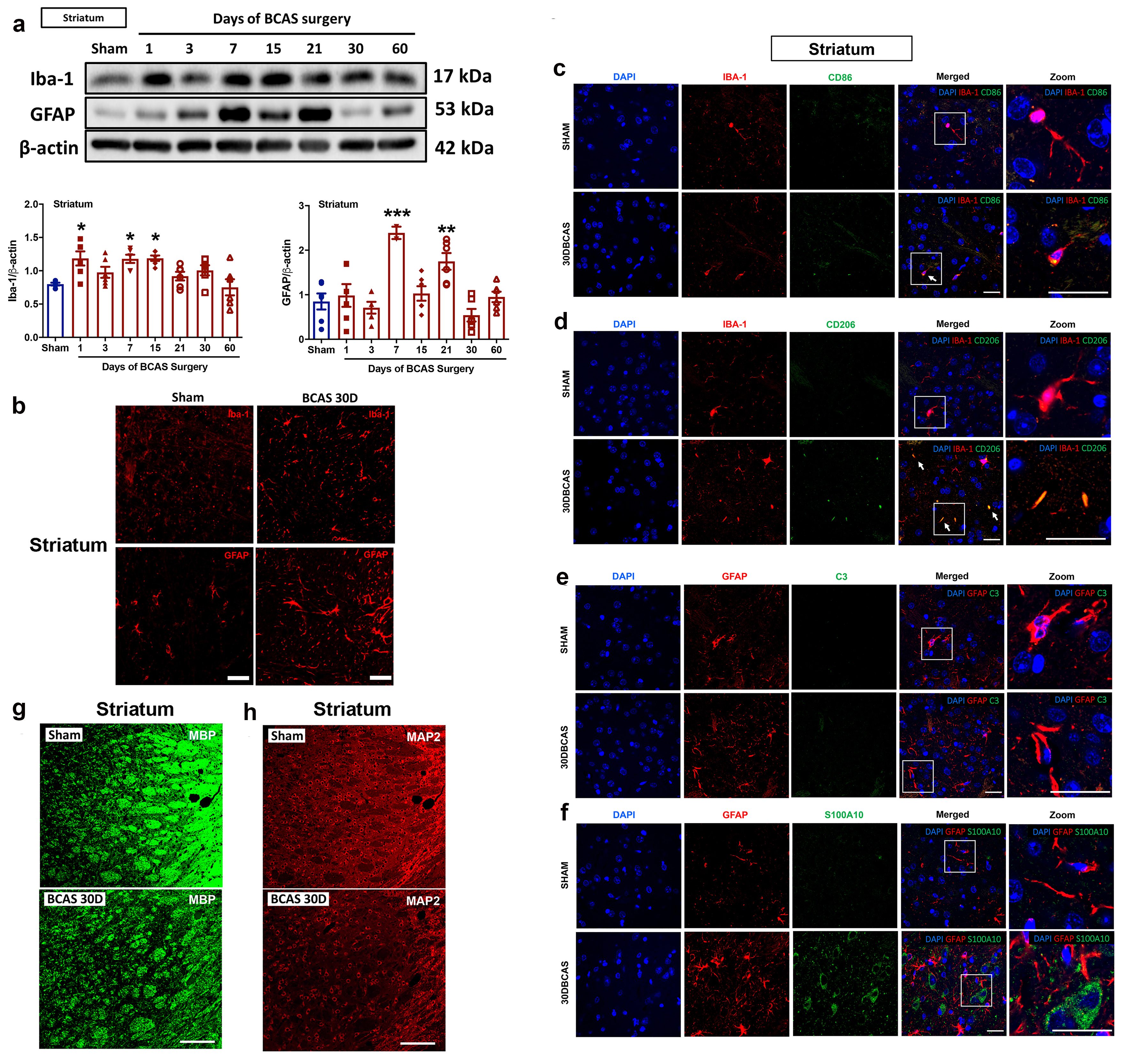

### S.Figure 11

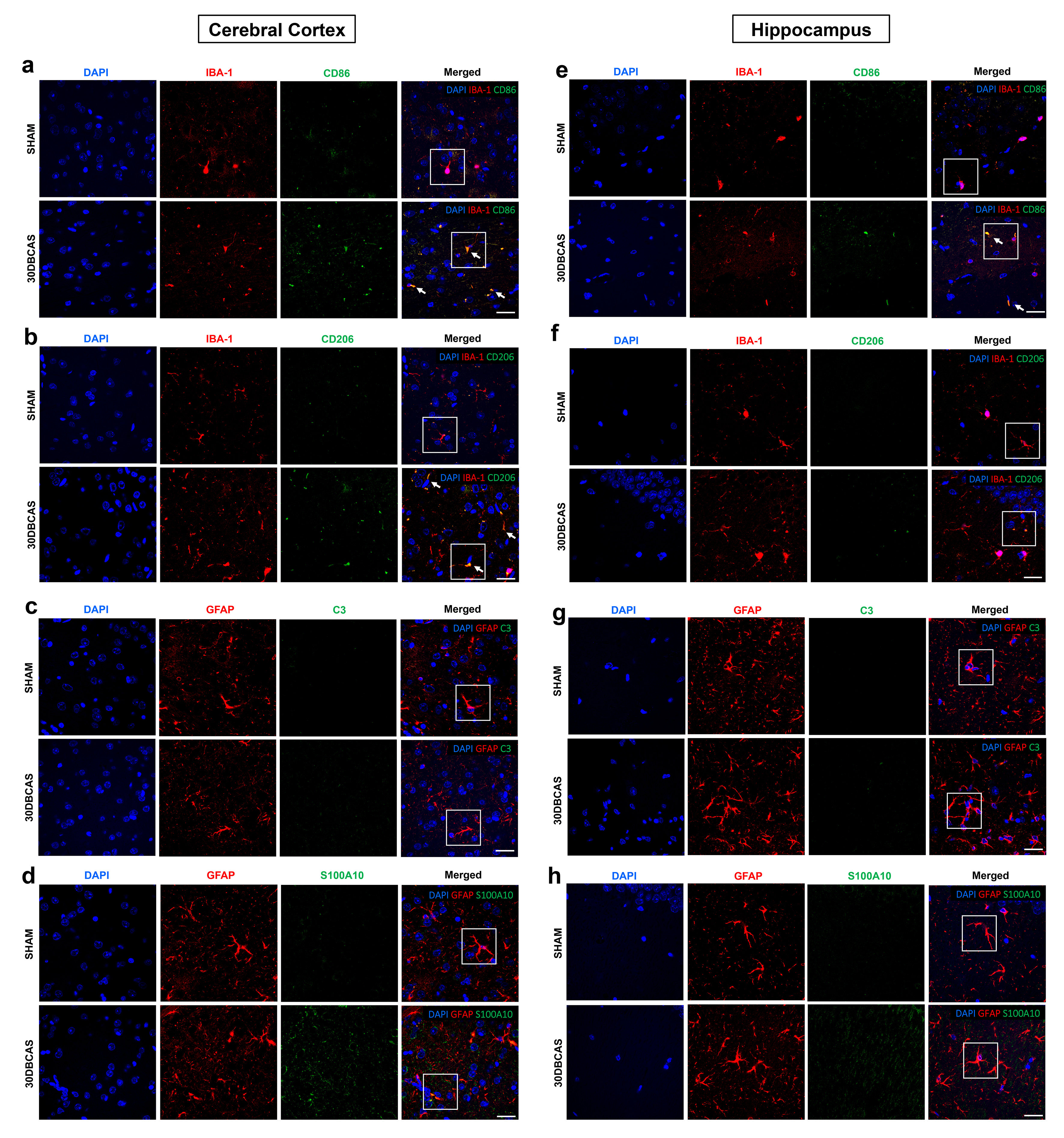

### S.Figure 12

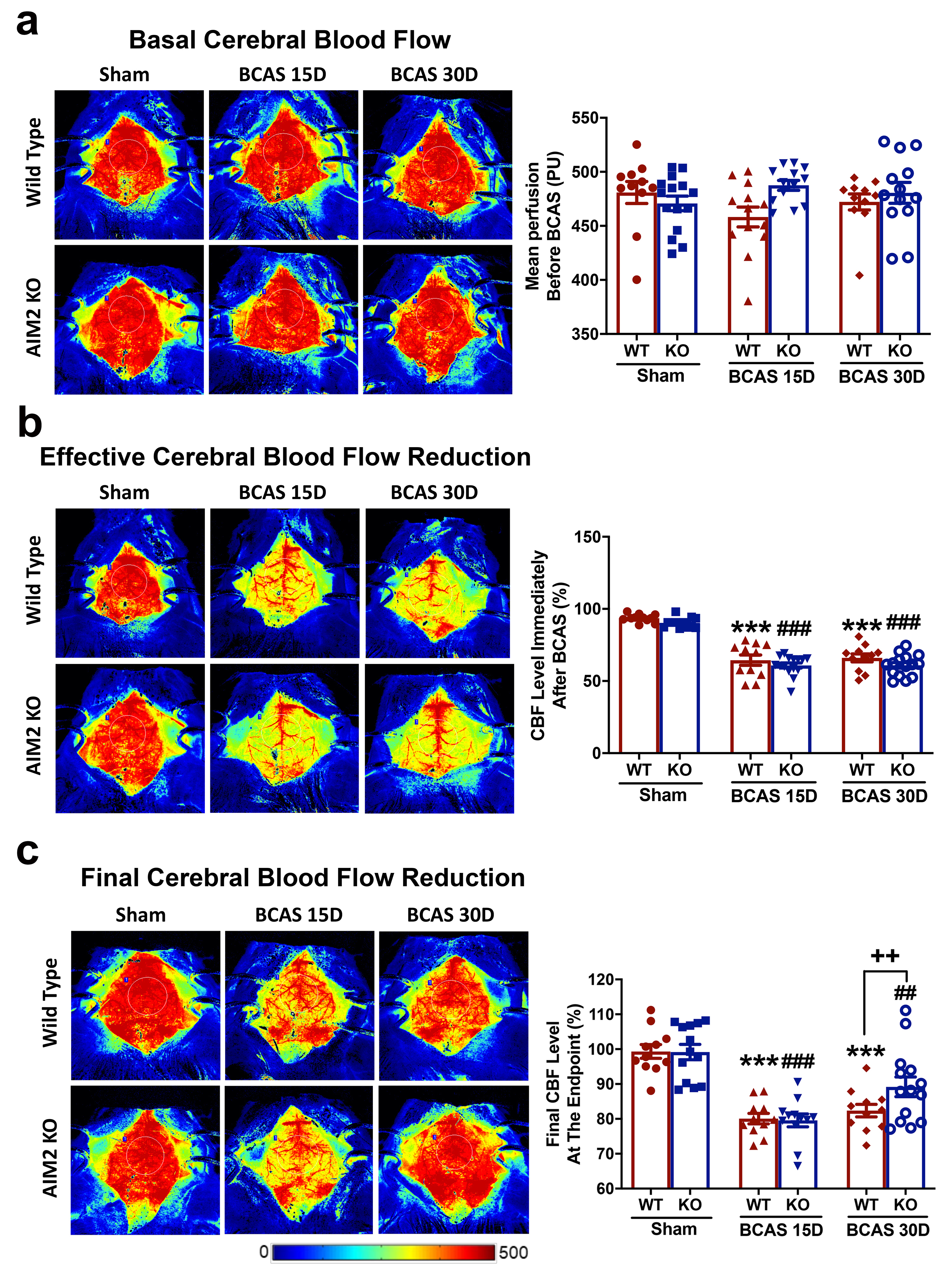

### S.Figure 13

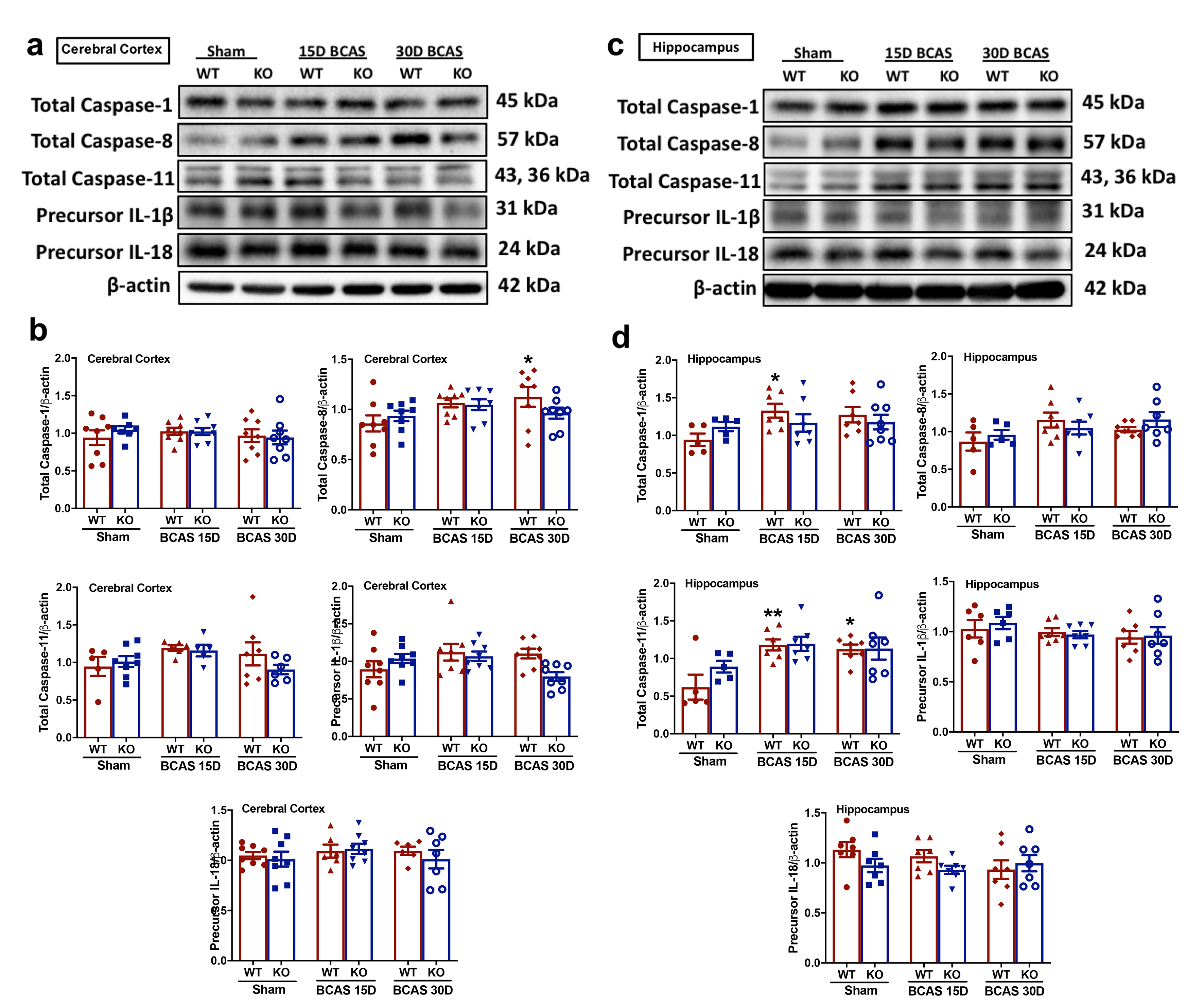

### S.Figure 14

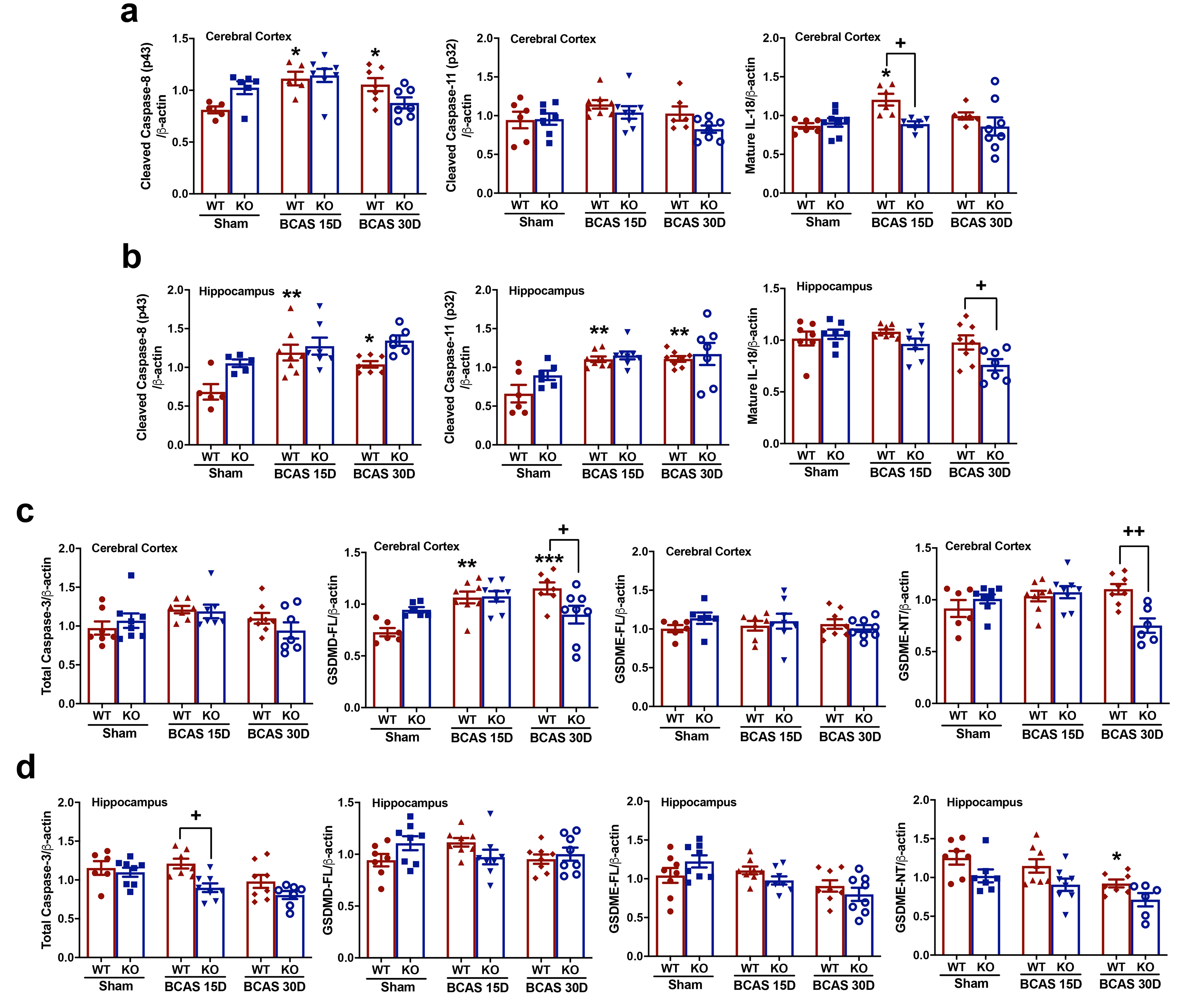

### S.Figure 15

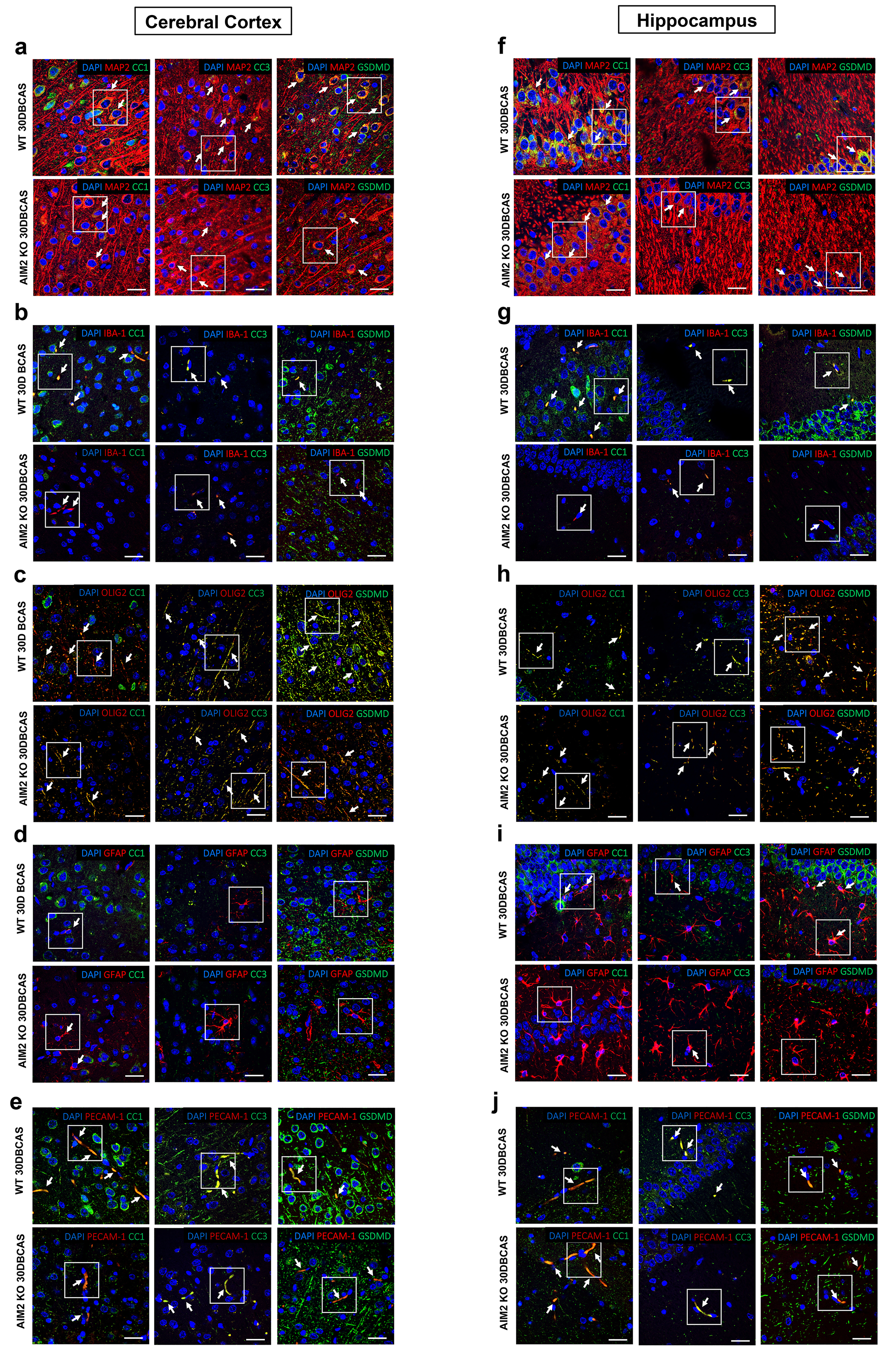

### S.Figure 16

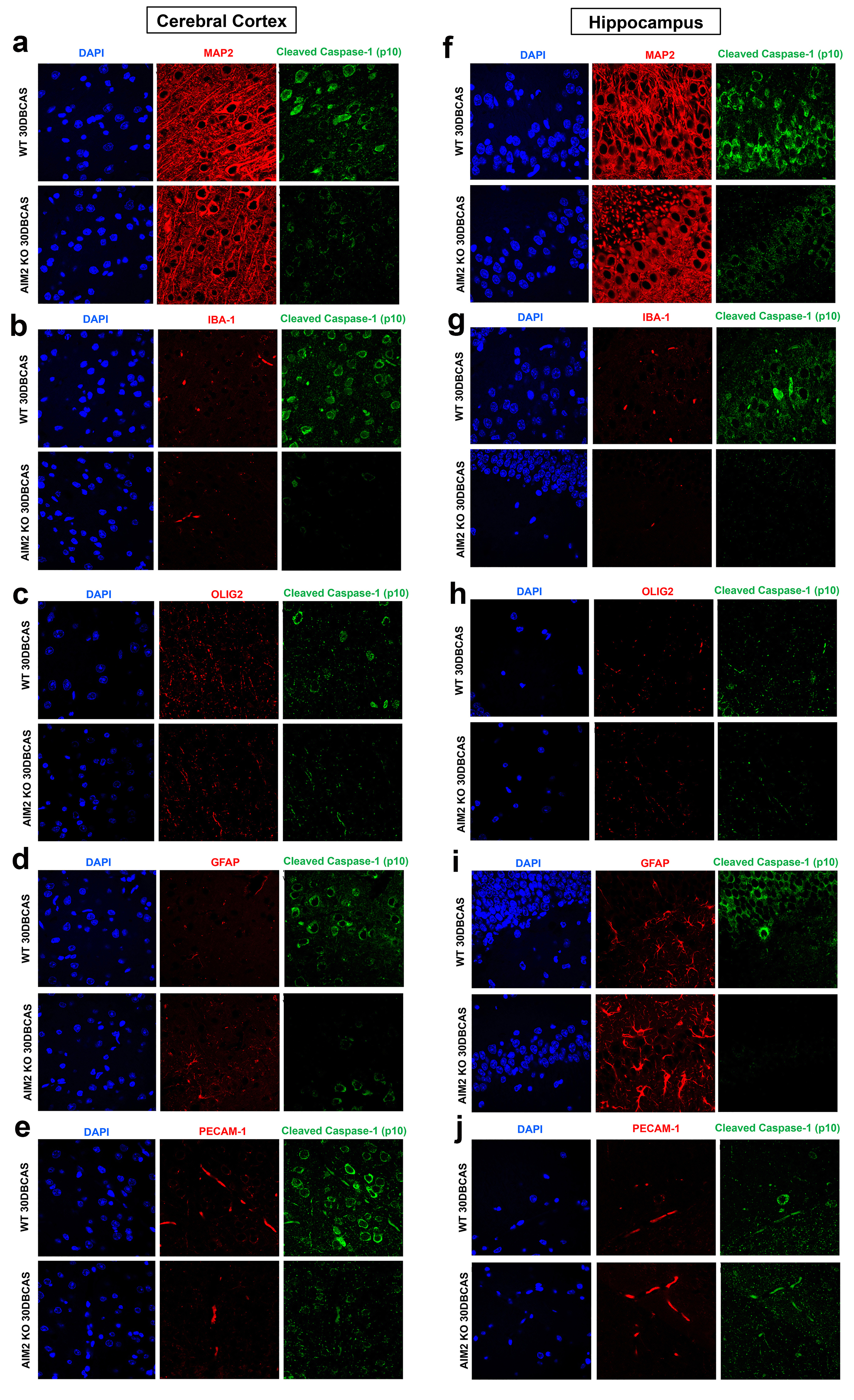

### S.Figure 17

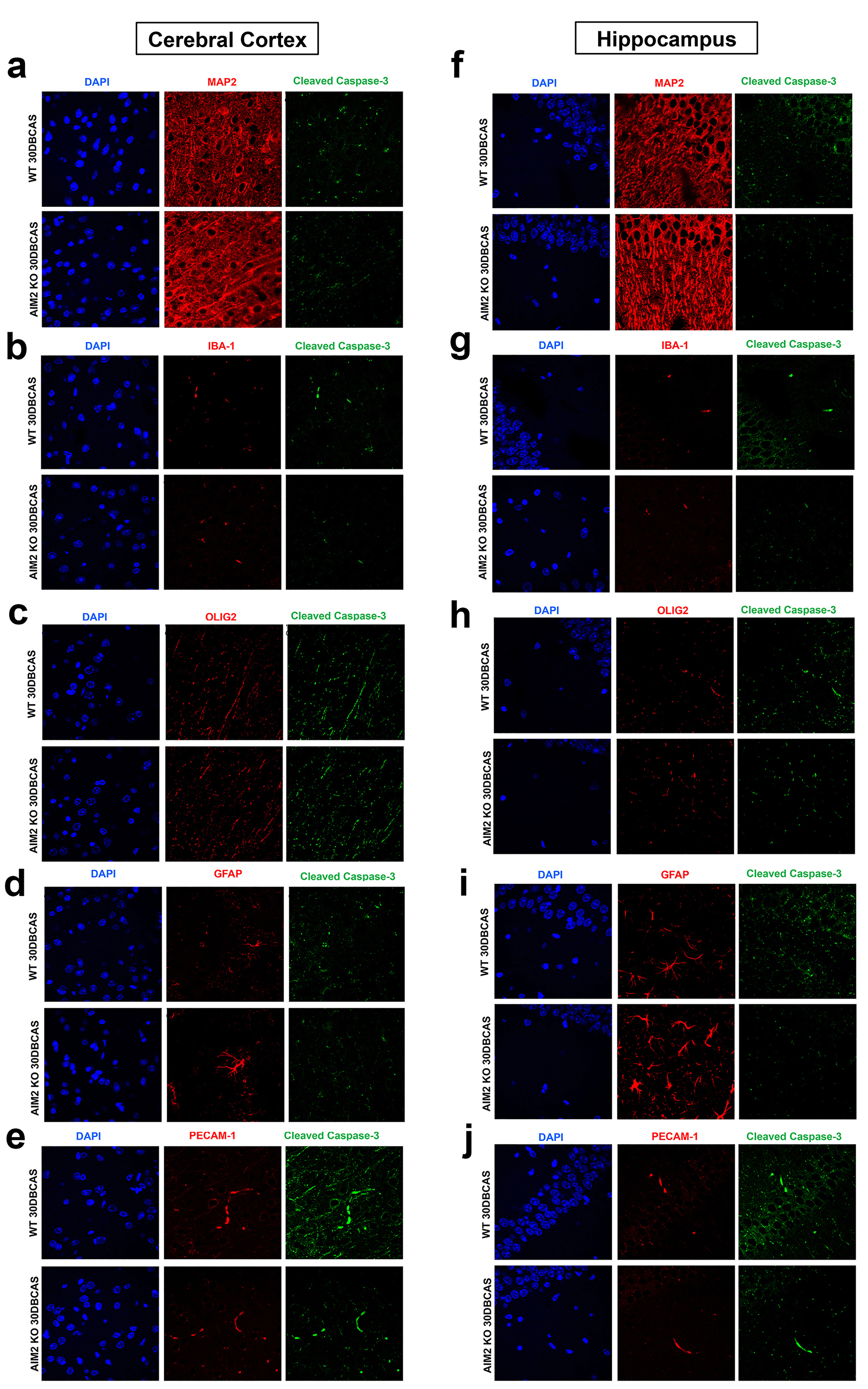

### S.Figure 18

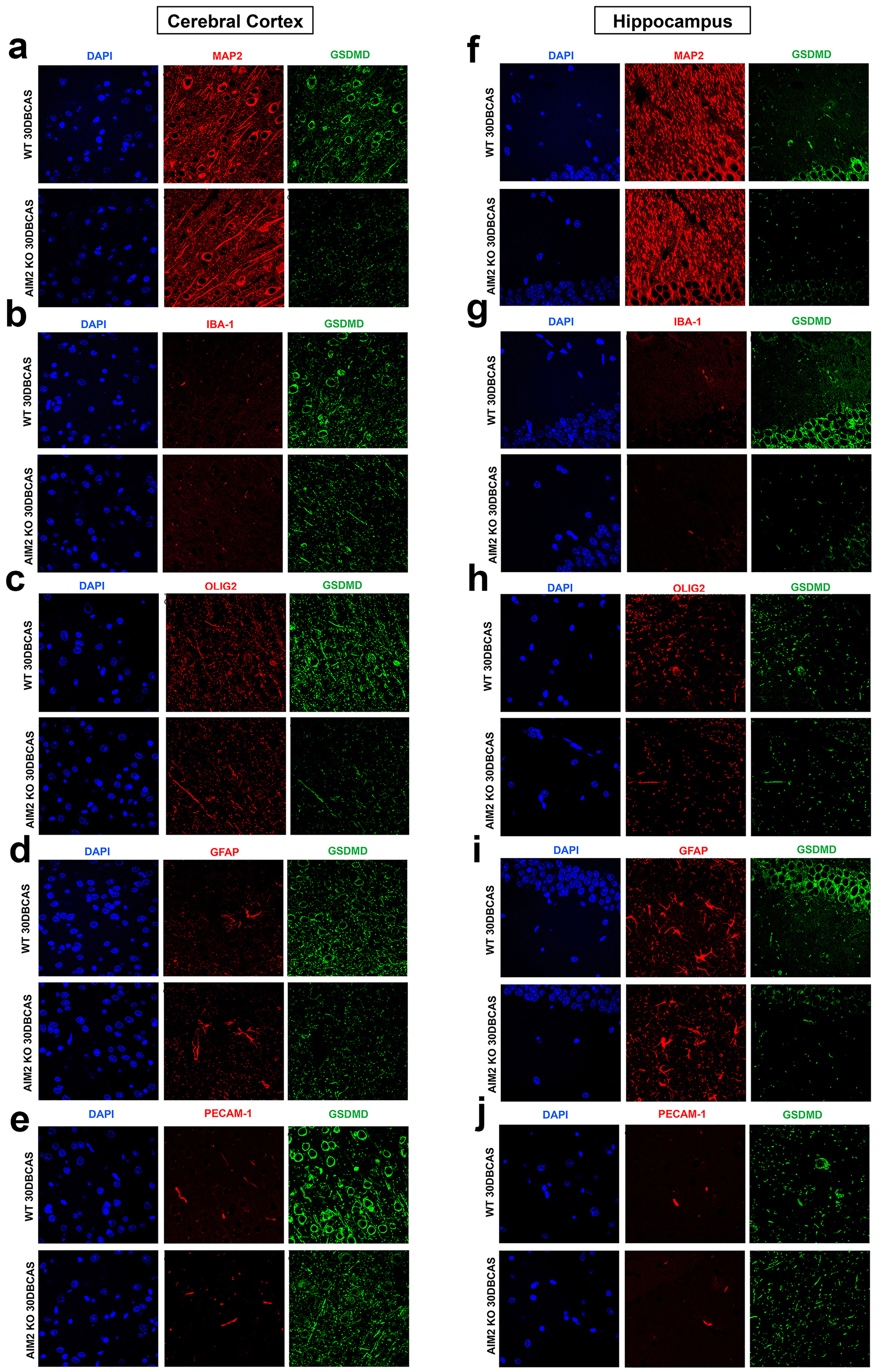

### S.Figure 19

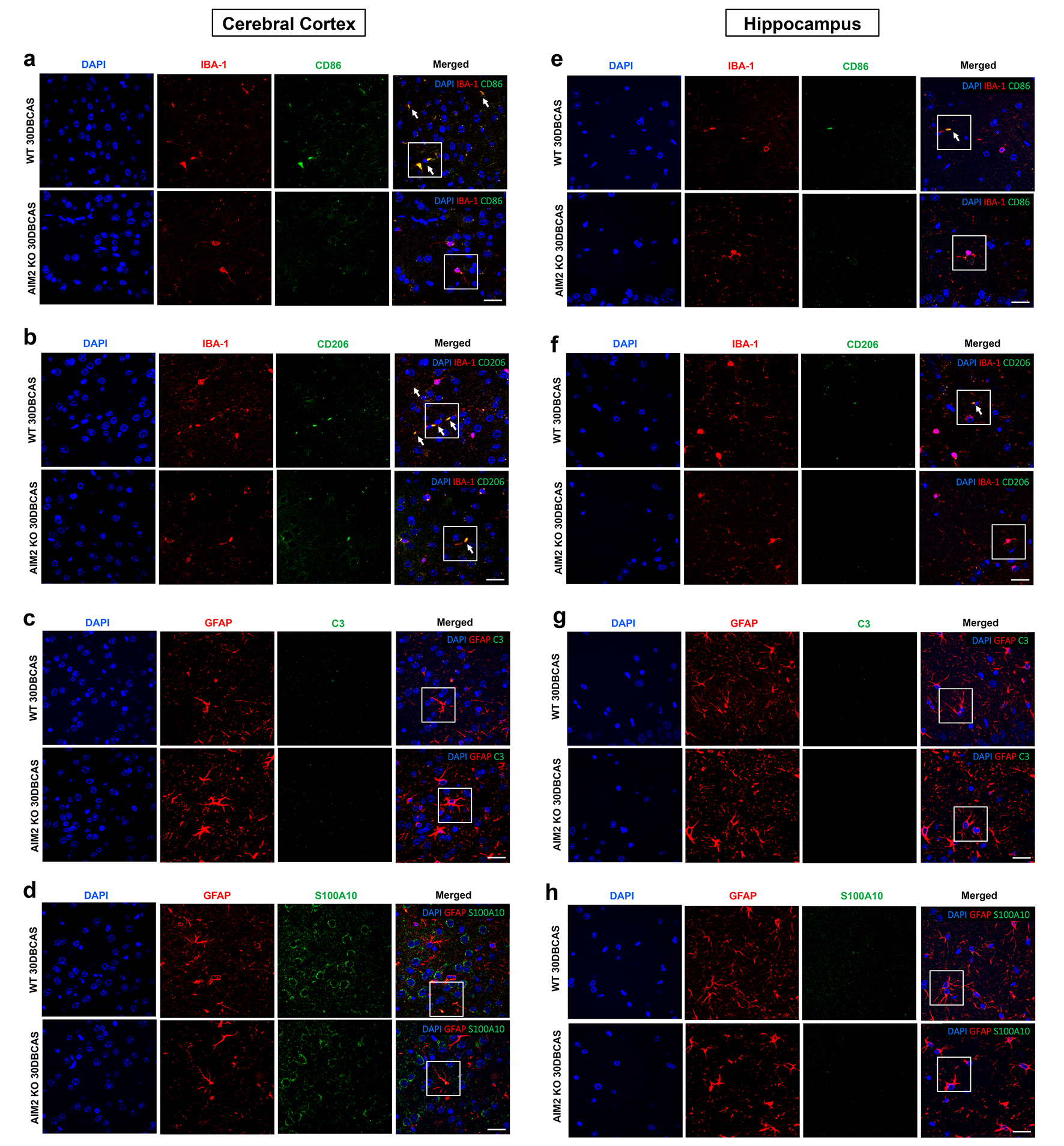
